# Identification of Novel Long non-coding RNAs and Expression Analyses in Rectal Neuroendocrine Tumors

**DOI:** 10.1101/2021.07.18.452620

**Authors:** Mahesh Kumar Padwal, Bhakti Basu

**Author notes:** **Corresponding author Dr. Bhakti Basu**, Molecular Biology Division, Bhabha Atomic Research Centre, Trombay, Mumbai – 400 085, India.

## Abstract

Altered expression of long-non-coding RNAs (lncRNAs) is associated with various pathological conditions, including cancers. LncRNAs associated with rectal neuroendocrine tumors (rNETs) remain unexplored. Here, we report the identification of 418 high confidence novel lncRNAs by analyzing publicly available RNA-Seq datasets of primary rNETs and rNETs with lymph node or liver metastases. Novel lncRNAs had basic characteristics similar to the known lncRNAs and showed individual-specific expressions. Expression analysis in the normal rectal mucosa and pancreas NETs (pNETs) and small intestine NETs (siNETs) revealed high expression of two lncRNAs, namely MSTRG30689.1 and MSTRG341161.1. Functional characterization revealed that 9 lncRNAs could work as the ce-RNA and one as miRNAs precursor. Cis- and trans- analysis of the 9 PCG-associated lncRNAs revealed that these lncRNAs could regulate various important cancer-related genes. Further, an integrated regulatory mechanism for selected cancer hallmark genes such as SMAD3, ZEB1, TCF4 and ATM through lncRNAs is predicted. Taken together, our analyses reveal common and unique lncRNAs associated with rNETs and their metastatic variants, their PCG targets, and predict mechanisms of lncRNA mediated regulation of target PCGs.

## 1. Introduction

Around, 90% of the human genome is transcribed into the RNA, however, only 2% of the transcribed genome code for proteins (Alexander et al. 2010; Pertea 2012). The remaining 98% of the RNAs, earlier believed to be transcriptional junk, is non-coding RNA(ncRNAs) (Ohno 1972). Biological functions of ncRNAs are emerging slowly and they are now being evaluated for potential biomarkers or therapeutic targets in various disorders including cancer (Watson et al. 2019). Based on their length ncRNAs are called small non-coding RNAs (sncRNAs) having a length of fewer than 200 bases or long non-coding RNAs (lncRNAs) which are longer than 200 bases (Djebali et al. 2012; Han Li and Chen 2015). ncRNAs perform either housekeeping functions through ribosomal RNAs (rRNAs) and transfer RNAs (tRNAs) or regulatory functions through microRNAs (miRNAs), small nuclear RNAs (snRNAs), piwi-interacting RNA (piRNAs), tRNA derived small RNAs (tsRNAs) and lncRNAs (Dozmorov et al. 2013). Further, based on regions on the genome where they are transcribed from, lncRNAs are classified as, intergenic, intronic, and antisense RNAs.

The regulatory functions of the miRNAs are very well studied in various types of cancers (Han Li and Chen 2015; Romano et al. 2017). Functional aspects of the lncRNAs are currently being explored. In breast, gastrointestinal, stromal, or prostate cancers, specific lncRNAs were shown to be involved in processes such as tumor suppression, alteration of the G1/S point, and growth enhancement (Fatima et al. 2015; Sanchez Calle et al. 2018). For example, metastasis-associated lung adenocarcinoma transcript 1 (MALAT1) was shown to play an important role in carcinogenesis by acting as a nuclear speckles scaffold for mRNA splicing in lung adenocarcinoma, breast cancer, and ovary cancer (Amodio et al. 2018). System-level Spatio-temporal expression profiling of the lncRNAs has been carried out in various cancers and diseases (Sanchez Calle et al. 2018). Johnson et al. have carried out the genome-wide transcriptome sequencing in the prostate cancer cohort and identified several un-annotated lncRNAs that were found to be associated with prostate cancer including PCAT1 which regulates cell proliferation (Prensner et al. 2011). Similarly, Ugne *et al*. have identified several lncRNAs deregulated in gastrointestinal stromal tumors (GISTs) and studied the role of the H19 and FENDRR in GISTs (Gyvyte et al. 2018). LncRNA either interact directly with target gene mRNA or bind to an RNA binding protein (RBPs) and direct it to mRNA (Zhang et al. 2018). In addition to these, the lncRNAs may act as miRNA precursors or miRNA targets (Romano et al. 2017). Functions of novel lncRNAs could be predicted by analyzing their co-expression with the protein-coding genes and their nearby gene loci (Cao et al. 2018).

Neuroendocrine tumors are tumors of the neuroendocrine system, derived from a special class of cells called neuroendocrine cells. Neuroendocrine cells are distributed throughout the body and have both endocrine and neuronal-like properties (Rindi and Wiedenmann 2011). According to the SEER data, NETs of the gastrointestinal tract (GI-NETs) account for approximately 58% of cases of all NETs and among the GI-NETs and rectal-NETs (rNETs) account for 20% (Man et al. 2018; Rodrigues et al. 2015). rNETs are further classified based on metastases to the regional lymph node or liver and around 75% of all GI-NETs metastases to the liver (Rodrigues et al. 2015). Global miRNAs profiling of small intestinal NETs by Su-Chen *et al* identified 5 up-regulated and 4 down-regulated miRNAs associated with tumor progression (Li et al. 2013). Similarly, several non-coding RNAs are identified in prostate NETs (Prensner et al. 2011; Zhang et al. 2018). However, expression profiling and functional relevance of lncRNAs in GI-NETs remain unexplored till date.

In this study, we have used a publicly available RNA-Seq dataset to identify novel lncRNAs associated with rNETs. We identified a total of 418 novel lncRNA transcriptscomprising 258 intergenic (lincRNA), 138 intronic, and 22 containing lncRNAs. The novel lncRNAs had lower transcript length and expression in comparison to the protein-coding genes with mean GC contents of 42.57%. Expression analysis revealed that some of the novel lncRNAs also had high expression in the siNET and pNET samples. A total of 2261 miRNAs were predicted to be targeting 418 novel lncRNAs. Next, we have carried out the functional characterization of these novel lncRNAs. Functional analysis of the lncRNAs showed that among novel lncRNAs 9 lncRNAs could work as ce-RNA, one as miRNA precursor and these lncRNAs may be involved in some of the important biological processes like epithelial cell proliferation, positive regulation of lymphocyte activation, protein kinase B signaling, adrenaline, noradrenaline mediated inhibition of the insulin secretion and activation of the GABA receptors. Further, various lncRNAs were found to be correlated with important cancer hallmark genes such ZEB1, SMAD4, KDR, TCF4, S100A8, S100A9 that are known to be involved in carcinogenesis. Finally, the possible regulatory mechanisms for some of the lncRNAs such as MSTRG11494, MSTRG10006, and MSTRG 33462, and MSTRG28734 with respect to the important cancer hallmark genes have been predicted.

## 2. Materials and Methods

### 2.1. Sequencing libraries and annotation data retrieval

Sequencing read archive (SRA) files for the rNETs, siNETs, pNETs and normal rectal mucosa were downloaded from NCBI SRA (GEO: GSM2627022, (GSE: GSE77635) (Alvarez et al. 2018). Genocode annotations (release 29) and human genome sequence primary assembly GRCh38.p12 were downloaded from the genecode website (https://www.gencodegenes.org/human/). The latest long-non coding RNA annotations were retrieved from the LNCipedia (Volders et al. 2019).

### 2.2. Read processing, alignment, and transcript assembly

The downloaded SRA files were converted to the fastq files using the split-3 legacy option of the parallel-fastq-dump tool which allows the use of multiple threads for faster file dumping [https://github.com/rvalieris/parallel-fastq-dump]. Sequencing read quality parameters such as the read-length, adapter contamination, and PCR artifacts were checked using the FASTQC. Poor quality reads were filtered out using the trimmomatic using the parameters (LEADING: 20, TRAILING: 20, SLIDING WINDOW:5:20, MINILEN:35) in paired-end mode (Bolger et al. 2014). Alignment of the reads to the human genome (hg38 p.12, Ensemble GTF (Version 95)) using the splice-aware aligner STAR (version 2.6.c) (Dobin et al. 2013). For alignment, the star-index files were generated using the default parameters, and then, alignment was carried using the parameters given in Additional File1.

### 2.3. Transcriptome assembly and quantification

Transcript assembly was carried out using the Stringtie, which is a super-fast, sensitive python-based transcript assembler (Pertea et al. 2015). The assembled transcriptome from the individual library was merged using the string-tie merge option with default parameters.

### 2.4. Identification of novel long non-coding RNA

The first step in the identification of long-non coding RNA is a comparison of the already known annotations with the de-novo assembled transcriptome. Merged transcriptome assembly was compared against known gene annotation from (Ensemble: Version 100, LNCipedia: Version 5.2, NCBI RefSeq annotation: (Release 109.20200522) using intersect bed tools (Neph et al. 2012). Any transcripts overlapped to any of the above assemblies filtered out using an in-house bash script. Further transcripts present on the alternate loci and patches filtered. Gffcompare was used for classifying transcripts based on their location in comparison to the protein-coding genes (PCGs) (Pertea and Pertea 2020). We have extracted three classes; (u) transcript of the unknown intergenic region, (c) transcript contained reference transcripts in its introns and, (i) transcripts with intronic regions. Transcripts with the exon count (>2), transcripts length (>200) were filtered using awk script. Filtered transcripts sequences were extracted from the genome region using the “gffread”. CPAT was run with the default parameter and transcripts coding score less than 0.38 and transcripts classified as non-protein-coding by PLEK were filtered out (Li et al. 2014; Wang et al. 2013). Six frame translation of the transcripts was carried out using the “TransDecoder” [https://github.com/TransDecoder/TransDecoder] and orf greater than 120 amino acid was filtered out us. Further PFAM search for all the remaining transcripts was carried out using hmmsearch against Pfam-A.hmm locally (Eddy 2011). Blat search was carried out using the human blat search on the UCSC blat web server.

### 2.5. Conservation score calculation

For conservation score calculation, phyloCons30way.wib file was downloaded from the USCS Table browser for the hg38 genome. The conservation score for the transcript in bed format was used for the extraction of the mean conservation score from the phastcons30way.wib file using hgWiggle tool of the UCSC “http://genome.ucsc.edu/admin/git.html”. The conservation at the gene level was calculated by taking the mean of each base.

### 2.6. Statistical analyses

Transcript length, GC contents and, conservation score comparison among protein-coding transcripts (PCTs) and known-lnc transcript and novel lnc-RNA transcripts were carried out as described previously(Kornienko et al. 2016). Briefly, the known lncRNA and PCTs were subsetted using the random function in r. The *p*-value was calculated thrice for each analysis using the Mann–Whitney U test and mean *p* values from three sets are reported.

### 2.7. Functional classification of the lncRNAs

For the *cis*-acting role of the transcripts the genes within the 100 kb region of the lncRNAs using the closet feature of bedops (Neph et al. 2012). For *trans*-acting roles of identified lncRNAs, Pearson correlation coefficients with PCGs were calculated using the python script provided with mec hRNA(Gawronski et al. 2018). For calculation of the correlation coefficient, counts for lncRNAs and PCGs were imported in R. In the first step, low expression genes were filtered out (Count Value < 10 in < 3 Sample). Next, the genes with very little variation in expression were filtered by taking upper quantile variant genes. Raw counts were normalized using the variance-stabilization transformation and PCC was calculated using the mechRNA provided scripts (Gawronski et al. 2018). For the ce-lncRNAs integrated network generation, first, the lncRNAs with more than 2 miRNAs binding sites were filtered, followed by the intersecting the PCGs which are positively correlated with these lncRNAs. The network was imported in the Cytoscape for visualization.

### 2.8. miRNA precursors and target prediction

lncRNAs that can be targeted by the miRNA were predicted using the miRANDA tool with a score and energy threshold of 160 and −15.0 kcal/mol respectively with strict seed matching at 5’ end to reduce the false-positive prediction (John et al. 2004). For miRNA precursors analysis miRNA precursors sequence were blasted against the lncRNA sequencing using local blast. lncRNA sequences that have significant hit (E10-6) were checked for the presence of the hair-pin loop using RNA fold. 2D structure of the lncRNA was drawn using a foRNA web-server.

### 2.9. Gene ontology (GO) enrichment analysis and sashimi plot

Gene enrichment analysis of the novel and differently expressed lncRNAs were carried out using the cluster profiler R package (Yu et al. 2012). Sashimi plot for the transcripts was plotted using the BAM coverage plotting tool “ggsashimi” (Garrido-Martin et al. 2018). Venn diagram was prepared using the online webserver [“http://bioinformatics.psb.ugent.be/webtools/Venn/”].

### 2.10. Prediction of regulatory mechanisms

For the prediction of the regulatory mechanism of the novel lncRNAs, we have used the mechRNA novel regulatory mechanism prediction pipeline (Gawronski et al. 2018). Initially, the lncRNA and mRNA interaction was predicted using Inta2RNA. The binding of the RBPs is predicted using the script provided with mechRNA. Inference of the mechanism was carried out using the mechRNA “infer.py” python script. For the mechanism inference amongst a set of transcripts for each gene, a transcript that is primary PCTs for that particular gene and part of the genecode basic annotation is selected. Mechanism prediction was carried out in the apriori mode.

## 3. Results

### 3.1. RNA-Seq data search and retrieval

For identification of lncRNAs in GI-NETs, the NCBI SRA database was searched RNA-SEQ data-sets for NETs, which have been sequenced in paired-end mode with a depth greater than 50 million reads. A study with a total of twenty RNA-SEQ libraries meeting the requisite sequencing parameters was retrieved (Alvarez et al. 2018), of which two libraries were removed due to a lack of replicates. These samples comprised of primary rNETs (n = 3), rNETs with lymph node metastases (n = 4), and rNETs with distant liver metastases (n = 11) (Table 1). The downloaded datasets have a total of 1984 million reads. After filtering the low-quality reads 1717 million clean reads were retained for further analyses (Table 1).

### 3.2. Identification of novel lncRNAs associated with rNETs

For the identification of the novel lncRNAs, we have followed the stringent pipeline which includes three steps for filtering and identification of the novel lncRNAs (Fig. 1a). In the first step, reads were aligned to the human genome GRCh38.p12 using the STAR aligner (Dobin et al. 2013). Around 90% of the total reads were aligned to the genome (Table 1). The aligned bam files were then used for the identification of the novel lncRNAs. A python-based framework string-tie was used for fast and accurate reference-guided transcript assembly, with default parameters (Pertea et al. 2015). Each assembled transcriptome had on average 1,57,000 transcripts (Table S1). The assembled transcriptome in general transfer format (gtf) file from each sample were then merged using string-tie to remove the redundant and spurious transcripts. Merged gtf file had a total of 10.6 % novel loci (2,54,997 transcripts in 60831 loci) in comparison to the reference assembly (2,25,969 in 56653 loci) (Fig. 1b). Three stringent steps were followed to distinguish novel transcripts from the known transcripts. The first step was the filtering of the known transcripts, including coding as well as non-coding RNA. For this purpose, the merged transcript assembly was compared with annotation from LNCipedia (version 5.2), Ensemble genes (version 100), and NCBI’s RefSeq gene annotation which also includes predicted transcripts, using “bedtools intersect”. Transcripts that have overlap with exons of any transcripts from three assemblies were filtered out. In addition to this, transcripts present on the patches or alternate loci on the chromosomes were also removed. After the first step, a total of the 3840 potentially novel transcripts were retained (Fig 1b). In the second step, three filters were applied which included, transcript lengths (> 200 bp), exon counts (> 2), and class filter (intergenic, intronic and contained), and 861 transcripts were retained for further processing. In the third step, the transcripts were assessed for their coding potential using the four tools CPAT (Wang et al. 2013), PLEK (Li et al. 2014), ORF prediction by Transdecoder, and homology search by PFAM. CPAT and PLEK are the two most commonly used coding potential assessment tools while PFAM translates the transcript sequences into all 6 frames and does a homology search for the presence of the known protein-coding domains. After filtering out the transcripts with coding potential, 584 transcripts remained. Finally, to remove the transcripts which may align to multiple loci on the genome, BLAT alignment was carried out against the human genome and 153 transcripts that mapped to multiple locations were filtered out. A total of the 418 transcripts which include 258 intergenic, 138 intronic, and 22 containing transcripts were identified as novel lncRNAs Fig. 1c. These novel lnctranscripts were transcribed from 331 loci on the genome. Novel lncRNA transcript sequences and gtf file are given in Additional Files 2 & 3, respectively.

**Fig. 1:**
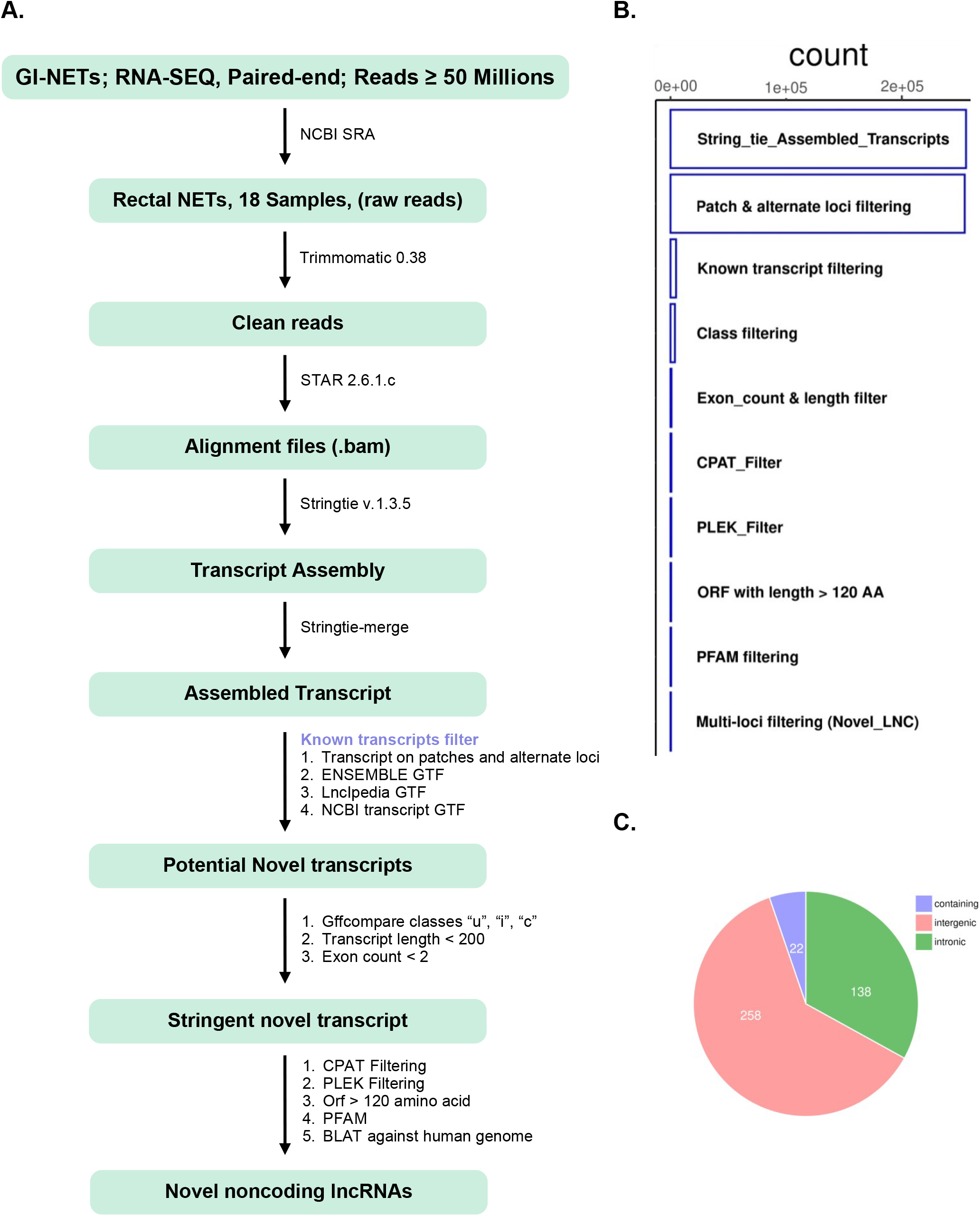
Stringent computational pipeline used for the identification of the novel lncRNAs. (a) Steps involved in the identification of the lncRNAs. (b) The number of the transcripts retained after filtering at each step. (c) Pie chart showing the distribution of the novel identified lncRNA in three classes intronic, intergenic, and containing.

### 3.3. Basic features of the novel lncRNAs associated withrNETs

Next, the novel lncRNAs were compared with the protein-coding and known lncRNA transcripts to evaluate their basic features such as transcript length, exon count per transcripts, percent GC content, and expression variability. First, we checked the distribution of novel transcripts on all the chromosomes (Fig. 2a and Table S2). Chromosome 4 transcribed the highest number of the lnc transcripts followed by chromosome 1 (Fig. 2a). Novel transcripts length ranged from 209 bp to 13 kb, with a mean length of 1728 bp. Unlike known lncRNAs, we found no significant difference between the length of the protein-coding transcripts (Mean 1831) and novel lncRNA transcripts (Mean 1728, *p-*value 0.31) (Fig. 2b). However, known lncRNAs transcript length (Mean 1514) was found to be significantly lower in comparison to the novel lncRNAs (Mean 1728, *p* value 1.936e^−05^). Novel lncRNAs contained fewer exons (median 2 exons per transcript) in comparison to protein-coding (median 6 exons, *p* value 2.2e^−16^), and known lncRNAs (median 3 exons, *p* value 2.2e^−16^) (Fig. 2c). The average GC content of the novel lncRNAs was 42.57 % which was lower in comparison to both protein-coding (Mean 51.35 %, *p* value 2.2e^−16^) and known lncRNAs (Mean 46.87, *p* value 2.2e^−16^) (Fig. 2d). We have checked for the conservation of the novel lncRNAs using the phastCons30way score (conservation score calculated by multiple alignments in 30 vertebrate species) for each base and then the mean score for each of the lncRNA transcripts was plotted. The novel lncRNAs had significantly lower conservation score (Mean 0.11) in comparison to the known lncRNAs (Mean 0.16, *p* value 0.004) and protein-coding transcripts (Mean 0.54, *p* value 2e^−16^) (Fig. 2e). Notably, 3 lncRNAs named MSTRG4415, MSTRG16489.5, and MSTRG16489.4 had a phastcons30way conservation score greater than 0.5 (Table S3) indicating that they may be more conserved.

**Fig. 2:**
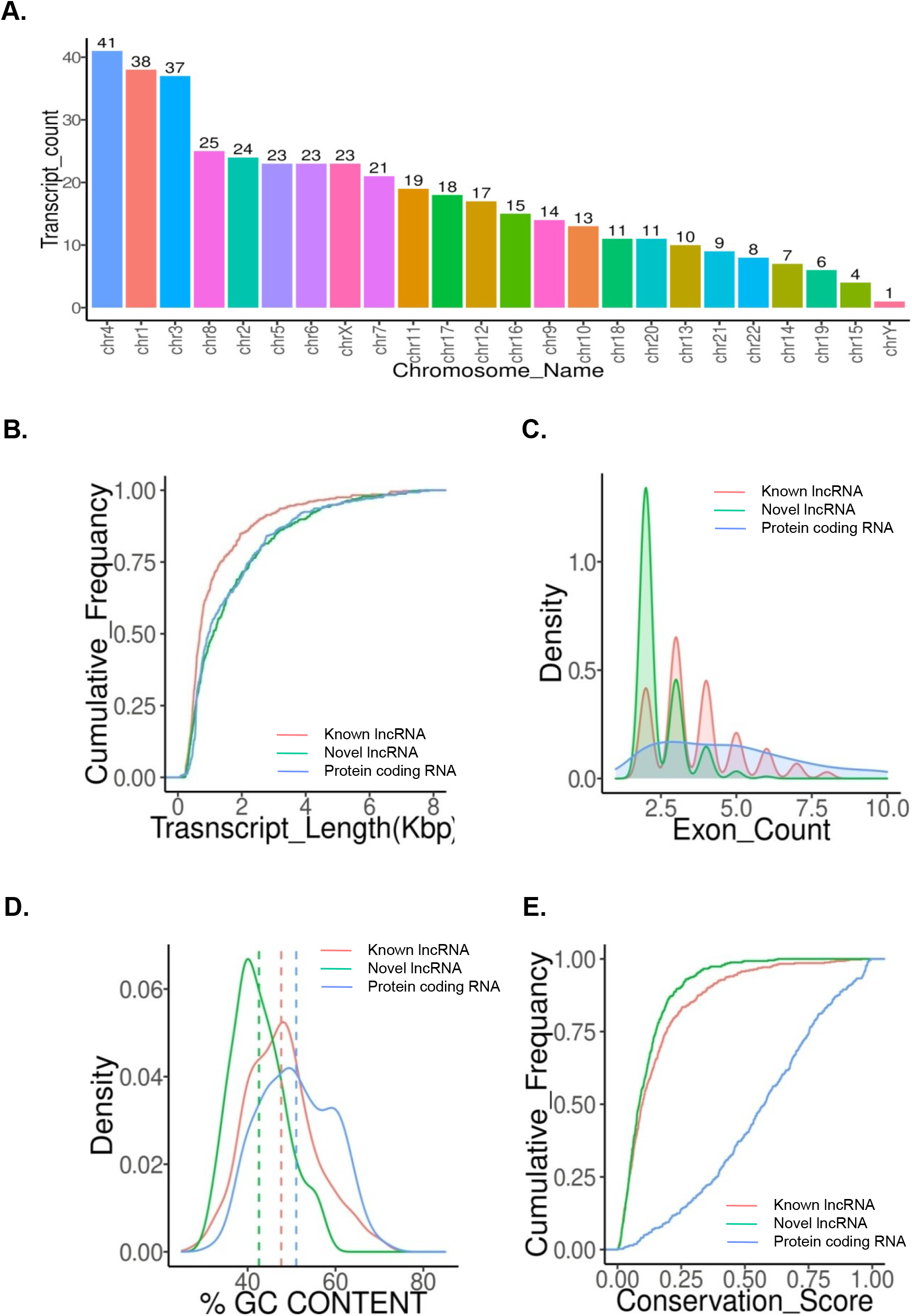
Basic characteristics of the identified novel lncRNAs in comparison to the known-lncRNAs and PCGs. (a) Chromosomal distribution of the identified novel lncRNAs. (b) Transcript length distribution of the identified lncRNAs. (c) GC content comparison of the novel, known lncRNAs, and protein-coding transcripts (PCGs). (d) Exons count comparison of the novel, known lncRNAs, and PCTs. (e) Phastcons conservation score of the known, novel lncRNAs and protein-coding transcripts.

### 3.4. Expression profiling of novel lncRNAs in rNETs versus normal rectal mucosa

Expression analysis showed that the novel lncRNAs had a higher median expression in comparison to known lncRNAs but a lower median expression than protein-coding transcripts in rNET samples (n = 18) (Fig. 3a). Hierarchical cluster analyses of the expression levels revealed that around 5%, 25% and 70% intronic lncRNAs showed very high, moderate and low expression, respectively, in rNET samples (n = 18) (Fig. 3b) while 35% and 65% lincRNA had a moderate and low expression, respectively, in rNET samples (n = 18) (Fig. 3c). Top highly expressed lncRNAs (TPM value > 5) include, MSTRG11494, MSTRG40589, MSTRG19919, MSTRG33062, etc. were found expressed in more than 70% of the samples (Table S4). Fifty percent of intronic and lincRNAs showed individual-specific expression, a similar observation has been reported earlier (Kornienko et al. 2016). Some of the novel lncRNAs showed a very high expression in a single sample, for example, MSTRG3010.2, MSTRG8058.1 (Figure S1a and S1b). When the log2(Mean_TPM) values were plotted against the coefficient of the variation (CoV) of the TPM, it showed that more than 50% lncRNAs had CV values greater than 2 suggesting a very high expression variability. However, the lower expression values could also possibly contribute to the observed expression variability.

**Fig. 3:**
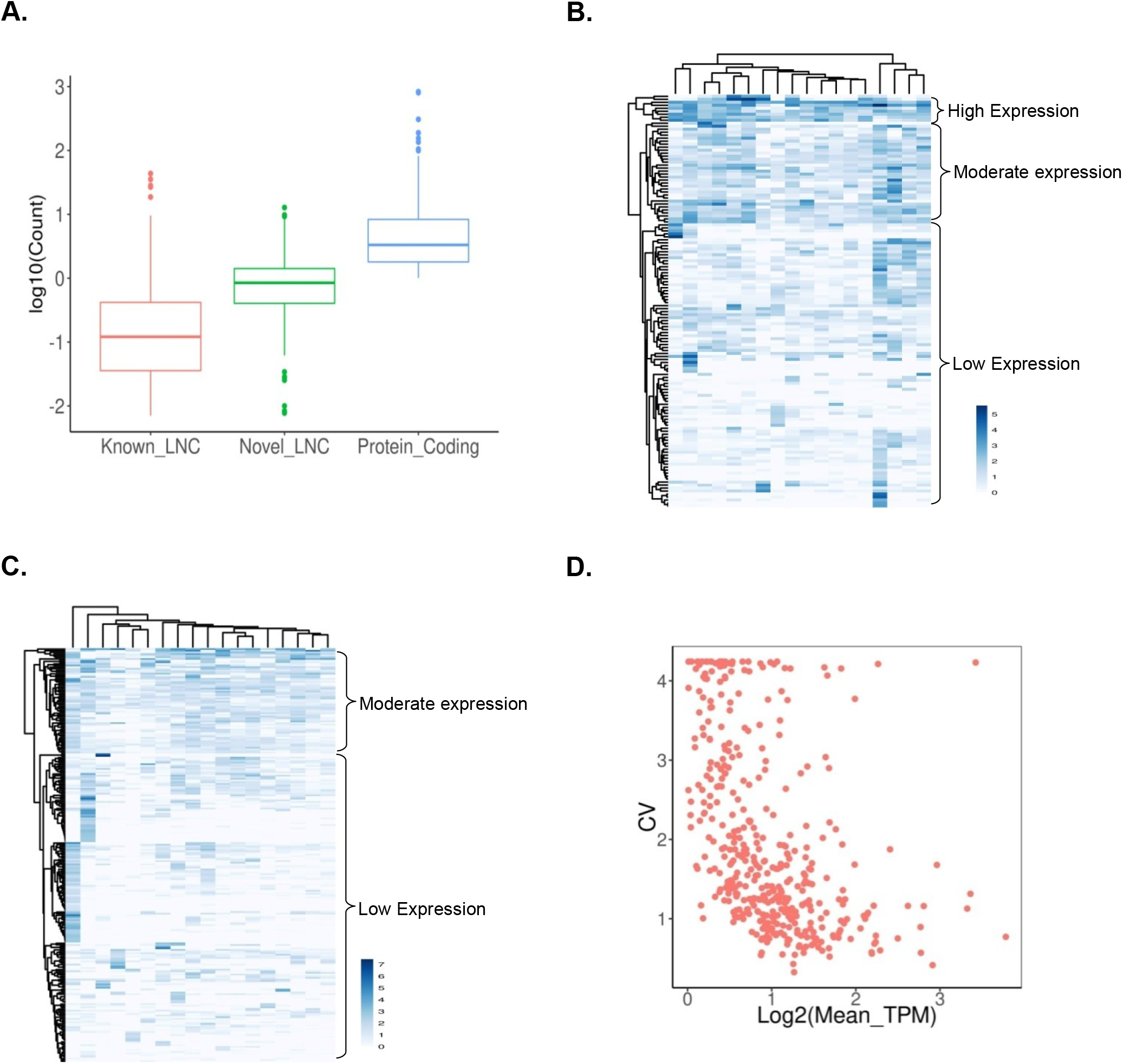
(a) Boxplot showing the comparison of the protein-coding transcripts, novel lncRNAs, and known lncRNAs. (b) Expression profile of the intronic lncRNAs. (c) Expression profile of the novel lincRNAs. (d) Scatterplot showing CV versus log_10_(TPM) values of the novel lncRNAs.

To check whether the novel lncRNAs identified in this study were specifically associated with rNETs, we checked if their expression was reported in the normal rectal mucosa. RNA-SEQ datasets for normal mucosa (GSE: GSE77635) (Hesson et al. 2016) were downloaded from the NCBI SRA database and analyzed. Out of 418 lncRNA transcripts, two transcripts named MSTRG30689.1 and MSTRG34161.1 showed a mean TPM value greater than 5 in normal rectal mucosa samples, 260 novel lncRNAs had no expression while the remaining showed moderate expression, which suggests that the majority of the novel lnc transcripts identified in this study are expressed only in the rNETs (Table S5).

### 3.5. Expression profiling of the novel lncRNA in other GI-NETs

We have analyzed the expression of the novel lncRNAs in 120 siNETs and 65 pNETs samples. RNA-SEQ datasets were downloaded from the NCBI SRA database (GEO: GSM2627022). Expression profiling of the novel lncRNAs in siNET and pNET is depicted in (Fig. 4a and 4b). For pNETs, 36 transcripts had mean TPM counts greater than one and five transcripts named MSTRG30689.1 MSTRG34161.1, MTRG33046.1, MSTRG31760.2, and MSTRG32922.1 had mean TPM expression greater than 5. MSTRG30689.1 showed the highest expression across all pNET samples (Table S6). Similarly, in siNETs 50 transcripts had a mean TPM count greater than 1 and, five transcripts MSTRG30689.1 MSTRG34161.1, MTRG33046.1, MSTRG31760.2 and MSTRG2952.1 had a mean TPM value greater than 5 (Table S7). Thus, two novel lncRNAs MSTRG30689.1 and MSTRG34161.1 showed high expression across all samples in both siNETs and pNETs.

**Fig. 4:**
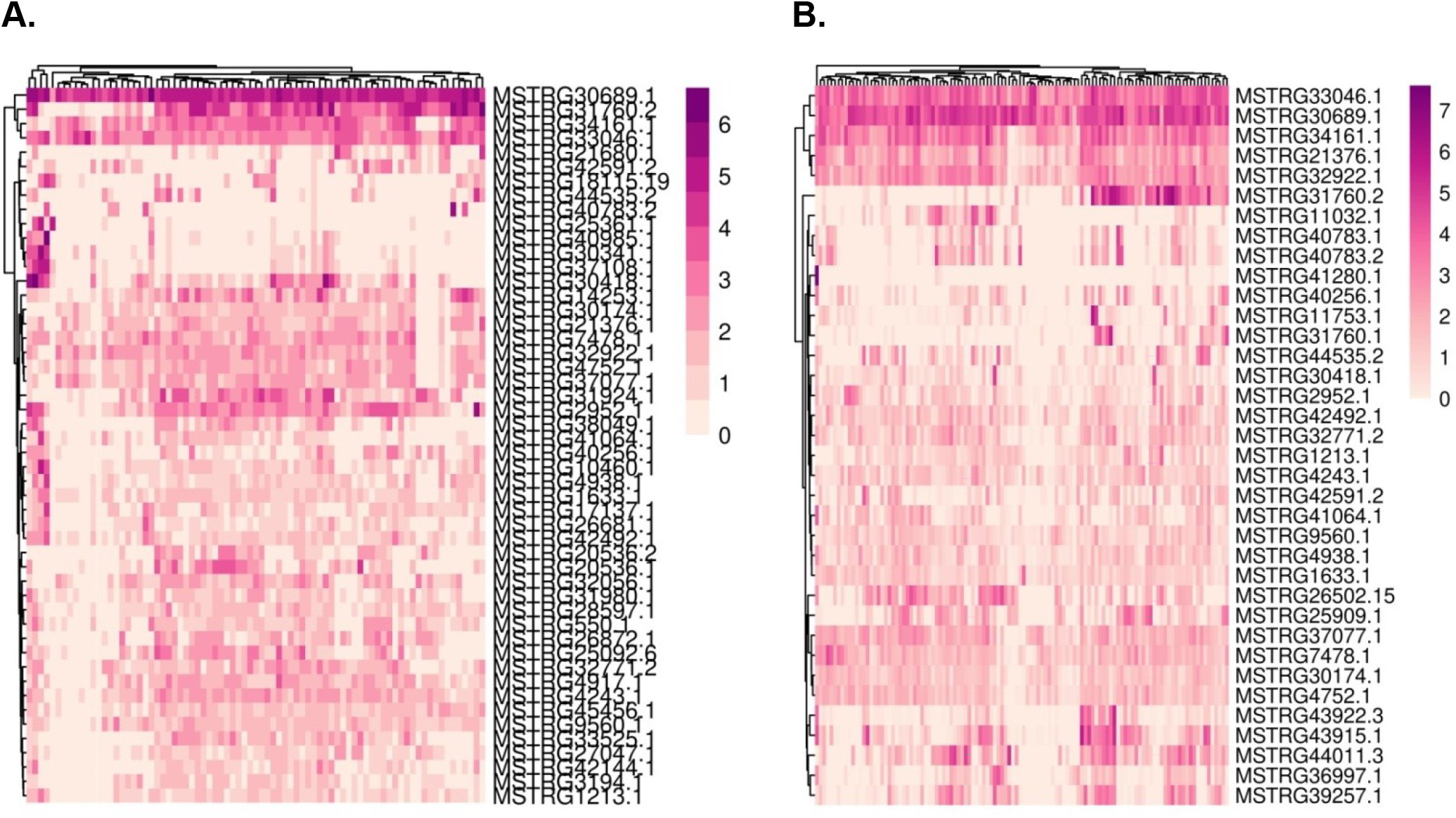
Expression profiling of the novel lncRNAs in (a) pNETs or (b) siNETs.

### 3.6 Localization of the novel lncRNA

The function of lncRNAs depend on their localization in a cell. The most probable location of the novel lncRNAs was predicted using a LocateR webserver (Ahmad et al. 2020). LocateR predicts the location of lncRNAs in the nucleus, ribosome, cytoplasm, or exosome. Among the 418 lncRNAs, 352 lncRNAs were predicted to be in the cytoplasm, 54 in the nucleus, 9 lncRNAs in the ribosome, and 3 lncRNAs were predicted to be localized in exosomes (Table S8).

### 3.7 Prediction of regulatory functions of the novel lncRNAs associated with rNETs

LncRNAs regulate gene expression at multiple levels by interacting through one or many components of the gene regulatory mechanism. For example, they interact directly with mRNA, act as a miRNA sponge, compete or help in the binding of the RNA binding proteins (RBPs), or act as chromatin modifiers(Sanchez Calle et al. 2018). To understand the functional contribution of the novel lncRNAs identified in this study, we checked for interactions of lncRNA with miRNA, *cis*- and *trans*-regulatory functions of lncRNA and lncRNA mediated RBP regulation and finally, we have integrated data from all regulatory mechanisms using the mechRNA to predict most probable regulatory mechanism.

#### 3.7.1. lncRNAs as ceRNAs

lncRNAs may work as competitive endogenous RNA (ceRNAs) wherein lncRNAs sequester miRNAs by competitive interactions through shared binding sequences resulting in altered expression of miRNA targeted genes (Cong et al. 2019; Wang et al. 2019; Zhou et al. 2019). To understand the role of the lncRNAs as ceRNAs, we used miRAnda tool (John et al. 2004) to analyze interactions between hsa-miRNAs and novel lncRNAs identified in this study. Our analysis revealed that a total of 2261 hsa-miRNAs could target all 418 lncRNAs (Table S9). The lncRNAs which act as ceRNAs have more than one binding site for miRNAs (Pellatt et al. 2016; Wang et al. 2019). In our analyses, a total of 28 lncRNAs had more than three binding sites for 36 hsa-miRNAs (Table S10). Strikingly, 17 hsa-miRNAs which were reported to be important in differentiating between normal rectal mucosa and rectal carcinoma (Pellatt et al. 2016) were found to be targeting the 88 novel lnc transcripts identified in this study (Table S11). Among these, two lncRNAs, MSTRG1633.1 and MSTRG31936.1 have three sites each for the three miRNAs (hsa-miR-17-5p, hsa-miR-20a-5p, and hsa-miR-93-5p) which have reported to be differentially expressed between the normal rectal mucosa and rectal carcinoma (Pellatt et al. 2016). To further confirm which lncRNAs with multiple-miRNA binding sites can work as ceRNA, we proceeded with the assumption that only the ceRNA localized in the cytosol could be positively correlated with miRNA-targeted PCGs. For this purpose, we first downloaded the experimentally validated miRNA-mRNA interactions from the miRTARBase database (Huang et al. 2020). Next, we filtered miRNAs that had more than 2 binding sites for lncRNAs and further, filtered miRNA-targeted-PCGs that were positively correlated with these lncRNAs (Fig. 5a). Finally, we created an integrated network of the miRNAs, positively correlated PCGs targeted by miRNA, and lncRNAs. The integrated network includes 9 lncRNAs, 19 miRNAs (Fig. 5b) and 48 PCGs sharing 109 interactions (Fig. 5c). Among these, MSTRG31936 is correlated with 27 PCGs and shared 6 miRNAs (Fig. 5c). MSTRG31936 regulated genes include various important transcription factors such as SOX4, Elk3, HOXB3, PRRX1, etc., all of which are shown to be involved in various types of cancers (Chen et al. 2016; Juang et al. 2016; Lee et al. 2017; Xu et al. 2018). MSTRG6729 is associated with 9 genes while MSTRG38037 associated gene includes FRMD3 which is experimentally proven tumor suppressor gene in lung cancer (Haase et al. 2007). In addition, when we checked the predicted location of lncRNAs, all were located in cytosol except MSTRG16391 which was ribosome-associated (Table S8). MSTRG16391 is associated with the two ribosomal associated genes RPL12 and RPL23 (Fig. 5c) which corroborated its location. The above examples clearly show that some of the novel lncRNAs, identified in this study, could play an important role in the regulation of the cancer-specific genes by acting as miRNA sponge.

**Fig. 5:**
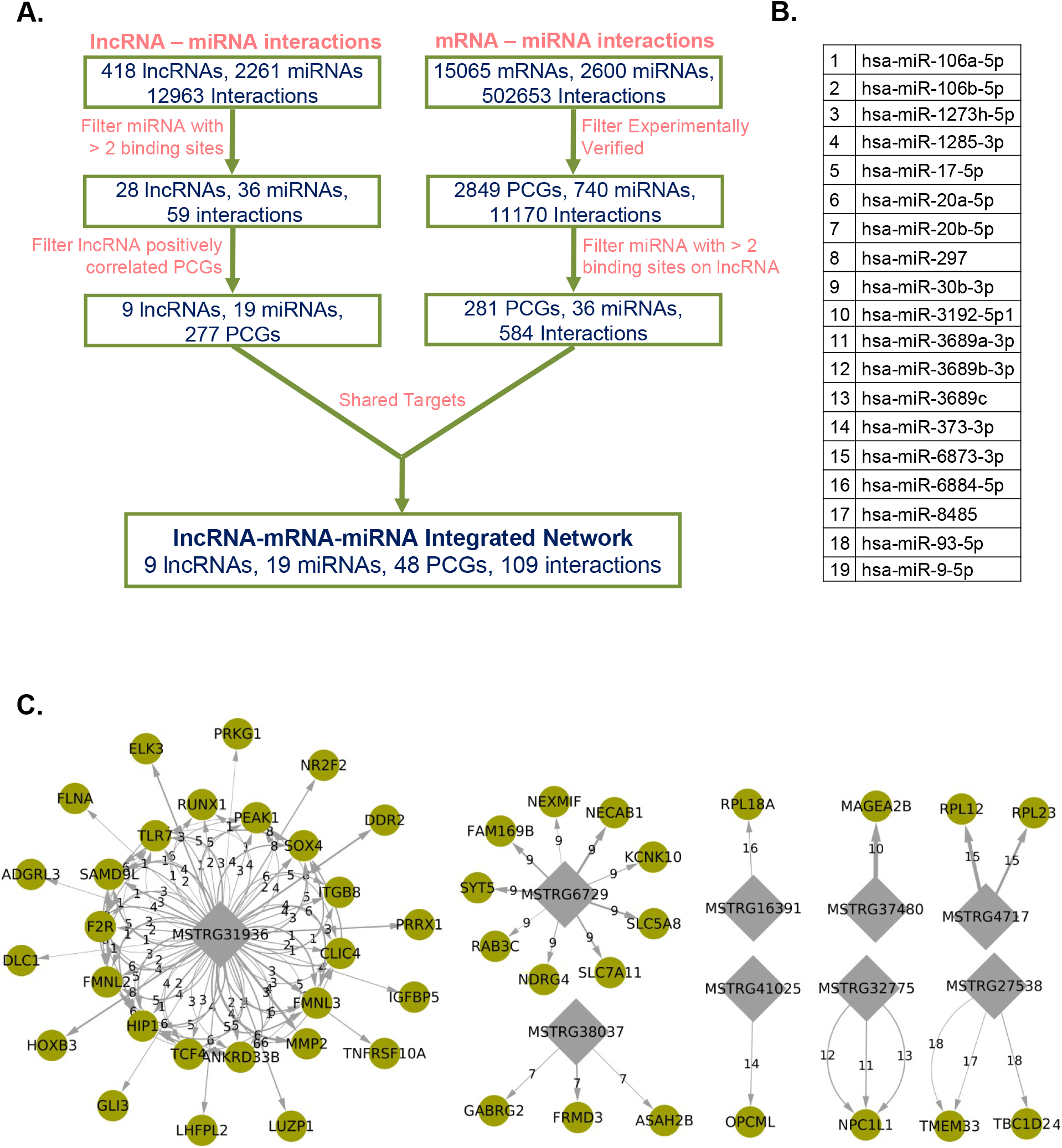
(a) Schematic representation for extracting functionally relevant protein-coding genes, miRNA associated with the ceRNAs. (b) The integrated network of the PCGs, miRNA, and lncRNAs visualized in Cytoscape.

#### 3.7.2. lncRNAs as miRNA precursors

For identification of lncRNAs that can act as miRNA precursors, first miRNA precursor sequences were blasted against the novel lncRNAs transcripts. Then the sequences which had an Evalue less than 10^−6^ were used for the prediction of the hairpin in the aligned region with mature miRNA sequence in the stem-loop. We found that lncRNA MSTRG2354.2 could act as a precursor for hsa-miR-619 miRNA. The structures of the hairpin loops of the precursor miRNA and the lncRNA are given in Supplementary Figure S2.

#### 3.7.3. *Cis* and *trans* regulation by the novel lncRNA

lncRNAs regulate the expression of nearby (*cis*-acting) or distant (*trans*-acting) protein-coding genes through various mechanisms (Chiu et al. 2018). For identifying *cis*-regulated genes, we searched for the genes which were located within 100 kbp upstream or downstream of the lncRNAs using BEDOPS closest feature (Neph et al. 2012). A total of 225 PCGs were identified in the vicinity of the 287 novel lncRNAs, which included 138 intronic lncRNAs (Table S12), 101 PCGs within 100 kb region of 127 linc transcripts (Table S13). Interestingly, 35 PCGs for 41 linc transcripts were located within 10 kb region (Table S14) while 7 bi-directional lincRNAs were located within 1 kb from 6 PCGs (Table S15). Some of these bi-directional lincRNAs are reportedly associated with the genes such as junction plakoglobin (JUP) gene which is overexpressed in oral squamous cell carcinoma (Fang et al. 2020), RAI2 which acts as the tumor-suppressor in colorectal carcinoma (Yan et al. 2018), MAP3K21 which increases migration and, MLN, GALK1, and TMEM151A are associated with metastasis in breast cancer (Marusiak et al. 2019; Williams et al. 2020).

To understand, the biological importance of these *cis*-PCGs, GO, and pathway enrichment analyses were carried out. GO analysis revealed that these *cis*-associated genes were specific to the cell-cell junction, an extrinsic component of the plasma membrane, G-protein beta/gamma subunit complex, adherence junction (Fig. 6a and Table S16). Reactome pathway analysis revealed that several pathways such as neuronal system, regulation of the insulin secretion, integration of the energy metabolism, cell-cell junction organization are found to be significantly enriched (Fig. 6b and Table S17). Some of the identified novel lnc transcripts were associated with a few pathways that are specific to the neuroendocrine systems.

**Fig. 6:**
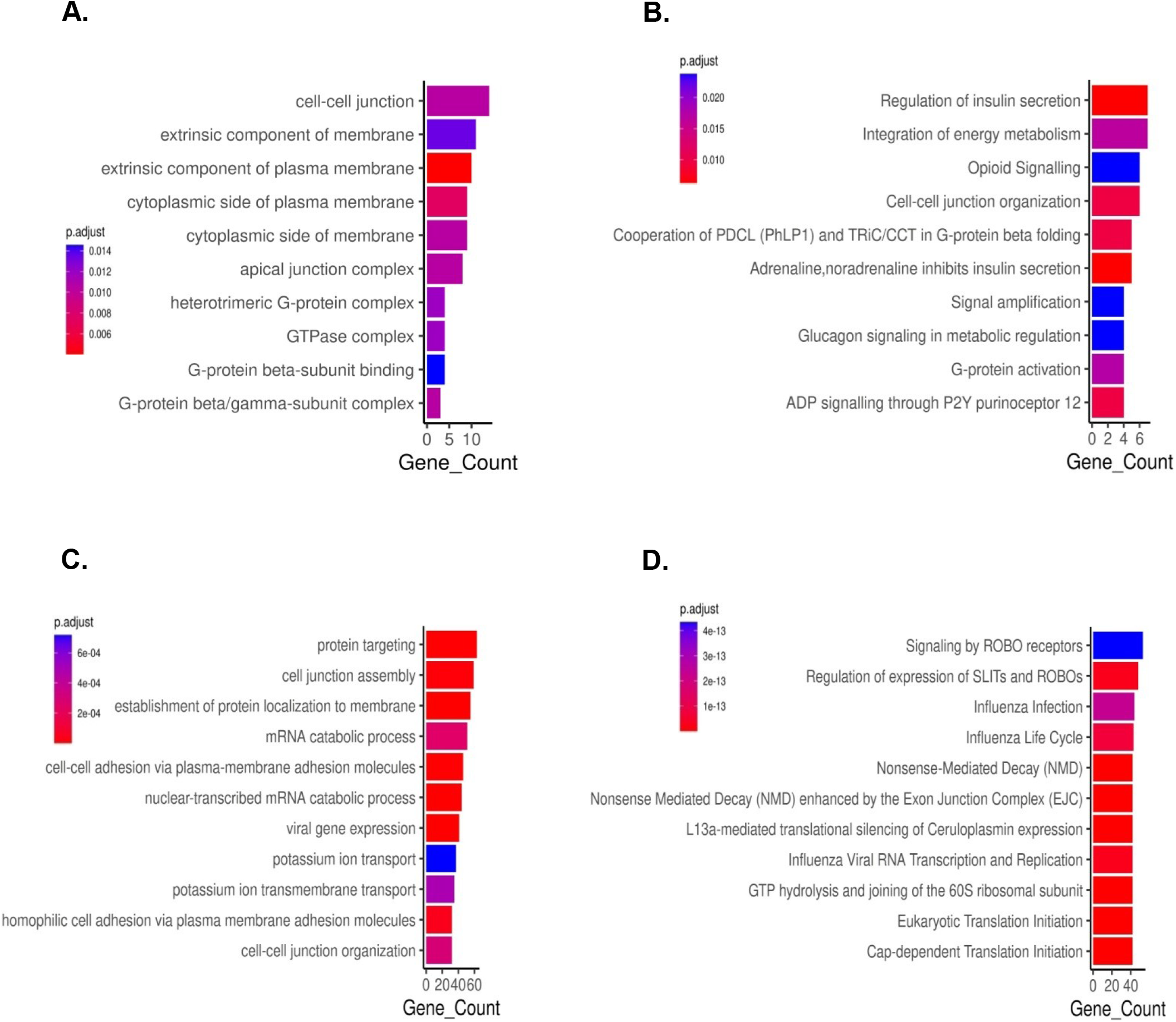
(a) Bar plot for significantly enriched GO terms (Cellular components) of *cis*-associated genes. (b) Bar plot for pathway enrichment analysis of the genes associated with lncRNAs in *cis*- manner. (c) Bar plot for significantly enriched GO terms (Biological processes) of *trans*-associated genes. (d) Bar plot for pathway enrichment analysis of the genes associated with lncRNAs in *trans*- manner.

To understand the *trans*-regulatory role of the novel lncRNAs, we have calculated the Pearson correlation coefficient (PCC) between PCGs and the novel lncRNAs. A total of 1445 PCGs were found to be significantly correlated with the 70 novel lncRNA genes (*p*.adjst ≤ 0.05) (Table S18). Pathway and gene enrichment analysis of these significantly correlated PCGs showed that the most overrepresented pathways are from the neuronal system, metabolism of the amino acids, ROBO receptor signaling, viral mRNA translation, influenza infection, etc. (Fig. 6c and Table S19). Similarly, the most important enriched biological processes were protein targeting, cell junction assembly, the establishment of protein localization in the membrane, synapse organization, neuron projection fasciculation, etc. (Fig. 6d and Table S20).

Further, we checked for the association of the novel lncRNAs with the cancer hallmark genes. For this purpose, we first downloaded a list of the cancer hallmark genes involved in more than 26 types of cancer (Nagy et al. 2021) and mapped these genes to the lncRNAs. Cytoscape networks for the cancer hallmark genes and lncRNAs are shown in Fig. 7. A total of 32 lncRNAs could be correlated with 40 cancer hallmark genes sharing 105 interaction. The cancer hallmark genes include some commonly misregulated transcription factors (ZEB1, SMAD2, REL, NR4A3, NFKBIZ), cell cycle-related genes (ATM, SMC1A, PRKDC, SMAD2, and SMAD4), oxidative phosphorylation related genes (NDUBF2, UQCR11, ATP5I) (Fig. 7). Further, some of the novel lncRNAs were correlated with more than 10 genes, for example, MSTRG33062 was correlated with a total of 12 hallmark genes (positively correlated with 5 genes, while negatively correlated with 7 genes). Interestingly, SMAD4 which is a tumor suppressor gene was negatively correlated with the 15 novel lncRNAs (Fig. 7) while it is known that SMAD4 deficiency promotes colorectal cancer progression (Papageorgis et al. 2011) Further, MSTRG16489 for which two transcripts, MSTRG16489.4 and MSTRG16489.5 are highly conserved (Table S3) was correlated with 30 PCGs (Table S21). Among its correlated genes include two important transcription factors SMAD4 and NCOA1 (Fig. 7). These examples underscore the importance of the identified novel lncRNAs in various carcinogenesis mechanisms.

**Fig. 7:**
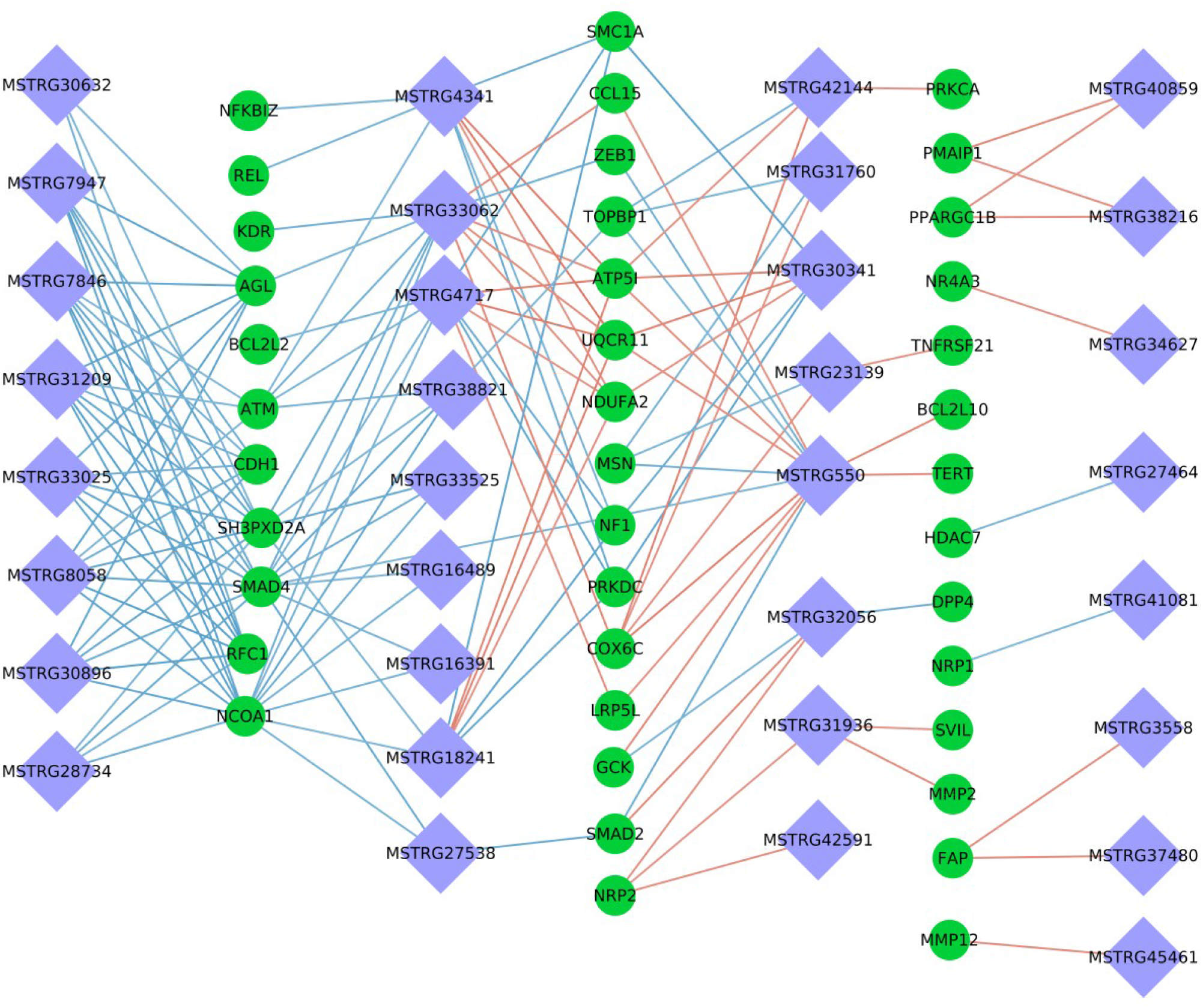
Association of lncRNAs with a set of cancer hallmark genes visualized in Cytoscape. lncRNAs (blue diamonds) which were positively correlated with the PCGs (green circles) are shown in red color, while lncRNAs which were negatively correlated with the PCGs are shown in blue color.

#### 3.7.4 RBP mediated regulation of the lncRNAs

RNA binding proteins are known to play an important role in the regulation of the lncRNAs and vice versa. For example, when bound to the lncRNAs ELVL1 allows their translocation from the nucleus to the cytoplasm (Briata and Gherzi 2020). Whereas, lncRNA TINCR is shown to help in localization of the IGF2BP2 on NAB2 gene and regulating the activity of the IGF2BP2 (Gawronski et al. 2018). To understand the regulatory role of the RBP associated novel lncRNAs, we used the mechRNAs RBP binding prediction machine-learning models to predict the binding of 22 RBPs to novel lncRNAs. These 22 RBPs regulate transcript stability, post-transcription regulation of target transcripts and translations. RBP binding peaks for the novel lncRNAs are given in Table S22. Cytoscape network for the RBPs and the lncRNAs is represented in Fig. 8. Seventeen lncRNAs were predicted to bind at least one RBP. Further, AGO1 was predicted to bind 313 lncRNAs (Fig 7, Table S22), while TARDBP was predicted to bind only 13 lncRNAs. The analysis showed that these lncRNAs could associate with several RBPs and may be involved in the regulation of various genes. Further, we have integrated these RBPs association to predict the mechanism of lncRNAs mediated regulations.

**Fig. 8:**
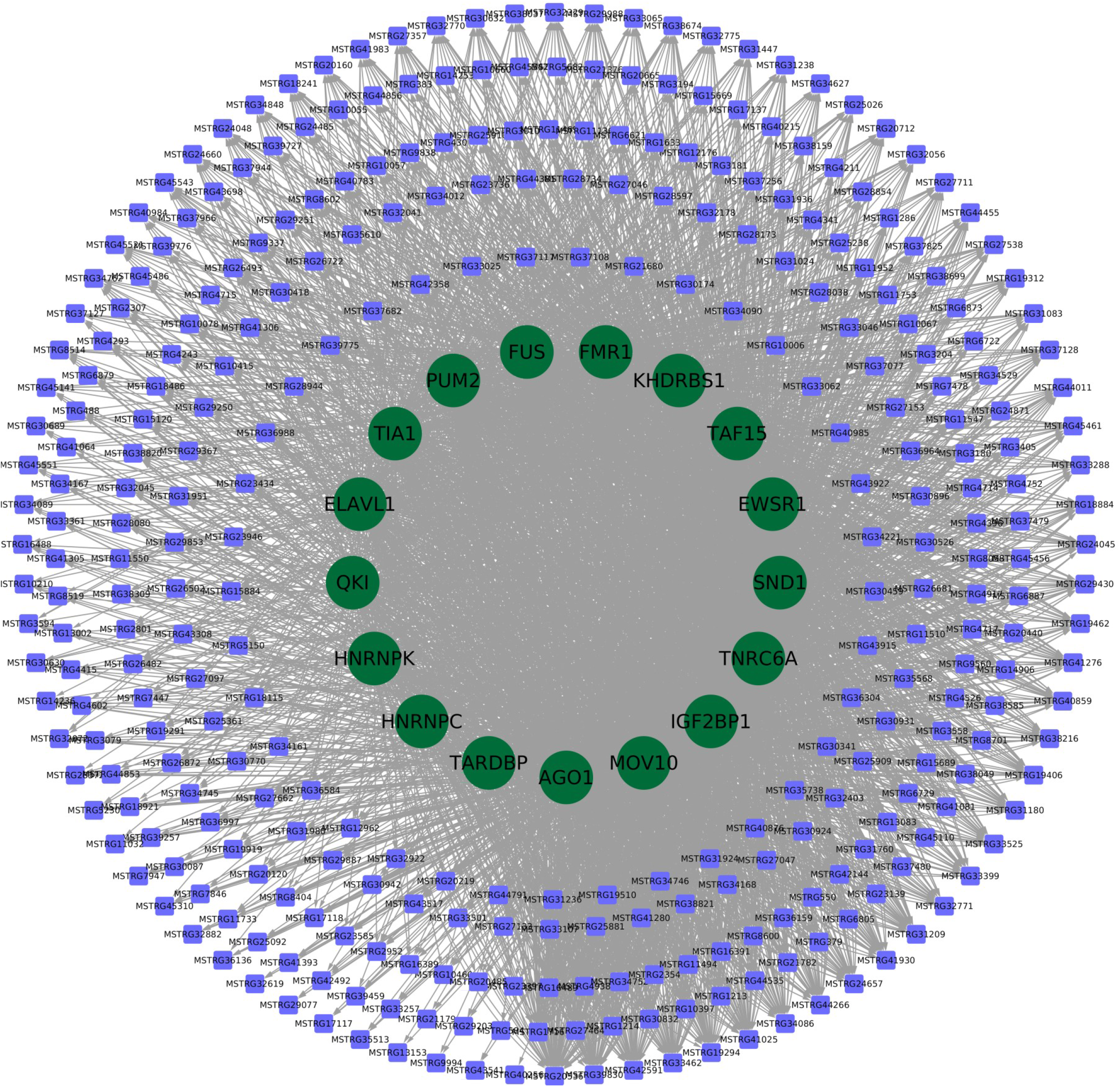
Association of the 17 RBPs (olive circles) with the novel lncRNAs (blue squares) visualized in Cytoscape.

### 3.8 Prediction of probable molecular mechanisms underlying lncRNA mediated regulation

To understand the regulatory mechanisms of important lncRNAs, we have used the mechRNAs pipeline which integrates data from mRNA-lncRNA interactions, RNA binding proteins (RBPs) mediated regulation, and lncRNA-mRNA correlation data to infer the most probable lncRNA regulatory mechanism (Gawronski et al. 2018). MechRNA allows the prediction of the mechanism at the transcriptome level, however, it requires high computation resources as well it predicts thousands of the probable targets for each lncRNA. MechRNA mechanism prediction can be run in two modes, apriori and without apriori information. In apriori mode, it considers the available information from the prior experiments and correlation coefficients provided to mechRNA. In without apriori mode, it considers the available information from, mRNA and lncRNA interaction, mRNA-RBP interaction, lncRNA-RBP interaction and predicts the final mechanism based on the consensus. We have run mechRNA in apriori mode to check whether the mechRNA could capture accurate information about the lncRNA-mRNA interactions. In case of ambiguity, we have used prior available information from experiments and correlated the data to find the most probable alternate mechanism for the lncRNA-mediated regulation. Since we were interested in understanding the role of novel lncRNAs in carcinogenesis, we have predicted the mechanism for the selected lncRNAs which were significantly correlated with some of the cancer hallmark genes or shown to play important roles in neuroendocrine carcinoma. Selected genes and their correlated lncRNAs include; BHLH4E1 and CRMP1 correlated with MSTRG11494; S100A8 and S100A9 correlated with MSTRG1006; ZEB1, TCF4, KDR, ATM and SMAD4 correlated with MSTRG33062; RFC1, SH3PXD2A and NCOA1 correlated with MSTRG28734. Further, the rationale for choosing these lncRNAs was (a) MSTRG11494 and MSTRG33062 were selectively and highly expressed in more than 70% of the rNET samples, (b) MSTRG10006 and MSTRG28734 were correlated with the important cancer hallmark genes that have important roles in tumor progression.

#### 3.8.1. MSTRG11494

MSTRG11494 is expressed from the locus on chromosome 13. It has two transcripts and MSTRG11494.1 showed higher expression in all samples of the rNETs (Supplemetnary Figure S3). MSTRG11494.1 transcript has a very low conservation score [0.054]. MSTRG11494.1 is an intergenic lncRNA and is positively correlated with the two important genes BHLHE41 and CRMP1. BHLHE41 is a basic helix-loop-helix transcription factor which is involved in EMT (Sato et al. 2016). It regulates the expression of the three master regulators of the EMT in neuroendocrine tumors, SNAI1, SNAI2, and TWIST1 (Sato et al. 2016). Similarly, CRMP1 plays an important role in axon growth, cell migration, and neuronal cone growth (Higurashi et al. 2012). In lung cancer CRMP1 has shown as anti-metastatic and anti-invasive (Tan et al. 2014). MSTRG11494 interacts with the BHLHE41 mRNA in the CDS region (Table S23). mechRNA predicted that MSRG11494.1 regulates BHLHE41 through direct upregulation via interaction at the CDS of the transcript (Combined *p* value < 0.0001) (Table 2). Similarly, for the CRMP1, direct up-regulation of the gene is predicted via RNA-RNA interaction at the CDS site (Table 2 and Table S23).

#### 3.8.2. MSTRG33062

MSTRG33062 is an intronic lncRNA, expressed from the intronic region of the gene Mastermind Like Transcriptional Coactivator 3 (MAML3), from a locus on chromosome 4. MSTRG33062 is correlated with 188 genes which include 108 negatively correlated and 80 positively correlated genes (Table S18). We have predicted the mechanism for the five important cancer hallmarks genes ZEB1, TCF4, KDR, SMAD4, and ATM (Table S23). ZEB1 and TCF4 play an important role in EMT (Zhang et al. 2019). The predicted mechanism for ZEB1 is through destabilization of the QKI binding to the ZEB1 (Table 2). QKI is shown to increase the stability of the ZEB1 (Han et al. 2019), and that is also visible in this dataset since QKI is positively correlated with the ZEB1 (Table S23). Similarly, for the TCF4, SMAD4, and ATM, de-stabilization of the QKI binding at the 3’UTR region is predicted. Interestingly, all four transcripts were positively correlated with the QKI. These show that the MSTRG33062 could target the QKI mediated stabilization of the transcripts. In addition, the free energy of the interaction between MSRG33062 and ATM is around −70 KJ, which denotes the MSTRG33062 could interact with the ATM with a very high affinity to replace the QKI. For KDR a direct down-regulation of the KDR through RNA-RNA interaction is predicted at the 3’UTR regions (Table 2).

#### 3.8.3. MSTRG10006

MSTRG10006 is expressed from a locus on chromosome 12. It has one transcript and it has been conserved more than the average score of novel lncRNAs (conservation score 0.14). MSTRG10006 is associated with two important genes named S100A8 and S100A9 (Table S23). S100A8 and S100A9 act as calcium sensors and they induce an inflammatory response in myeloid cells and other cancer cells (Srikrishna 2012). Both proteins were shown to be upregulated in various cancers such as pancreatic ductal carcinoma, breast cancer, prostate adenocarcinoma (Bergenfelz et al. 2015; Grebhardt et al. 2014). MSTRG1006 is positively regulated with both the proteins. For both S100A8 and S100A9, the direct up-regulation of both the transcripts via the RNA-RNA interaction is predicted (Table S23). For S100A8 lncRNA interaction is predicted at the 3’UTR region, while for the S100A9 it’s at the 5’UTR region.

#### 3.8.4. MSTRG28734

MSTRG28734 is an intergenic lncRNA, has two transcripts and, is expressed from a locus on chromosome 22. MSTRG28734 is correlated with 56 genes in *trans*. Three cancer hallmark genes RFC1, NCOA1, and SH3PXD2A are negatively correlated with the MSTRG28734. For RFC1 the predicted mechanism is the de-stabilization of FUS binding (Table 2). For SH3PXD2A also destabilization mechanism is predicted for the binding of the FUS protein (Table 2). FUS protein increases mRNA stability. For NCOA1, direct down-regulation of the NCOA1 via RNA-RNA interaction is predicted. The interaction range and site involved for interaction for all three lncRNAs and PCGs is shown in Table S23.

## 4. Discussion

The relevance of lncRNAs, one of the most common but poorly understood RNA species, to human diseases, is increasingly being realized. The promise of greater depths in RNA sequencing by the latest technologies has brought us closer to appreciating a regulatory link between lncRNAs and PCGs as a possible cause of several pathologies. Currently, genome-wide annotations and expression analyses of the lncRNAs are available for several cancers (Fatima et al. 2015; Sanchez Calle et al. 2018; Wang et al. 2019). Evaluation of quantitative associations between lncRNAs and cancer-related genes is being explored using computational methods and experimental evaluations (Cong et al. 2019; Prensner et al. 2011; Wang et al. 2019). Through these analyses, lncRNAs are implicated in crucial cellular processes such as apoptosis, autophagy, cell differentiation, cell proliferation, metabolism, etc. The lncRNAs regulate complex cellular mechanisms through (a) disrupting miRNA-mRNA homeostasis by sequestering miRNAs (Cong et al. 2019), (b) regulating translation or mRNA stability by base-pairing (Zhang et al. 2018), (c) epigenetically modulating gene expression by recruitment of chromatin-modifying factors (Amodio et al. 2018), etc. Thus, dysregulation in lncRNA could directly result in dysfunction in cellular homeostasis leading to pathophysiology.

Annotations and expression profiling of lncRNAs in GI-NETs remain completely unexplored. Here, our analyses on publicly available RNA-SEQ data-sets of primary rNETs and rNETs with lymph node or liver metastases, lead to the identification of 418 high confidence novel lncRNAs. These novel lncRNAs, belonging to 317 loci, were further classified into 258 lincRNAs, 138 intronic, and 22 containing lncRNAs. Novel lncRNAs showed standard features of lncRNAs such as fewer exons, lower transcripts length, less conservation, lower median expression and lower GC contents in comparison to the protein-coding transcripts. However, novel lncRNAs showed higher median expression than known lncRNAs. The majority of 418 novel lncRNAs showed low expression in the normal rectal mucosa, siNETs and pNETs. More importantly, the majority of the lncRNAs have individual specific expression, an observation that confirms that this can be a general feature of the lncRNAs. The phastCons conservation analysis revealed that three lncRNAs MSTRG16489, MSTRG4415, and MSTRG26502 were highly conserved.

Since lncRNAs could regulate mRNAs through altering the abundance or activity of miRNAs, we performed computational analyses to identify potential lncRNAs that could work as the ceRNA by integrating data from experimentally validated PCG-miRNA interactions and correlation data from our analysis. We have deciphered a set of 8 lncRNAs that could work as ceRNAs. PCGs targeted by these ce-lncRNAs include various genes, for example, Elk3, HOXB3, SOX4 which play important roles in the various stages of carcinogenesis. Further, our miRNA precursors identification analysis revealed that lncRNA MSTRG2354.2 could act as the precursor for hsa-miR-619 miRNA. Cis analysis of these lncRNAs with PCGs revealed that these lncRNAs are associated with various important cellular processes such as epithelial cell proliferation, mesenchymal cell proliferation, protein kinase B signaling, T cell activation, and differentiation etc. Cis-regulated genes include various important genes such as SMAD3, TGFBR2 and PTEN have significant associations with NETs. SMAD3 and TGFBR2 are downstream targets of tumor suppressor protein encoded by Multiple endocrine neoplasia type 1 (MEN1) where MEN1 modulates the transcriptional activity of these transcription factors (Kaji et al. 2001). PTEN expression variations are associated with grades and Ki-67 index in pancreatic NETs (pNETs) (Han et al. 2013). Further, trans association analysis of the novel lncRNAs with PCGs revealed that these lncRNAs could play an important role in carcinogenesis-related pathways such as cell-cell junction organization, regulation of expression of the SLIT and ROBO receptors. Besides, several neuroendocrine system-specific processes such as the regulation of insulin secretion, G protein activation, and various genes related to the neuronal system such as KCNN1, CAKM2D, and NRG1 etc. were computationally predicted to be associated with novel lncRNAs. NRG1 is a ligand for the EGFR and it plays an important role in the differentiation and growth of epithelial, neuronal, and glial cells (Jeong et al. 2014). Our trans-correlated PCGs include a set of the 40 cancer-hallmark genes, which shows that the identified lncRNAs could be regulating these important genes and vice-versa. Finally, we have predicted the mechanism for the cancer hallmark genes regulation by the lncRNAs. Our analysis has revealed that MSTRG33062 interacts with QKI binding site in four important cancer hallmark genes SMAD4, ATM, ZEB1, and TCF4. QKI has already been proven to play an important role in the EMT by stabilizing the ZEB1 mRNA and MSTRG33062 could regulate the stability of the ZEB1 and thus EMT.

Taken together, we report annotations of a set of novel lncRNAs, associated with rectal neuroendocrine tumors and their metastatic variants, their PCG targets, and predict mechanisms of lncRNA-mediated regulation of target PCGs.

## 5. Conclusions

We have reported the discovery of 418 novel lncRNAs which could enrich a growing list of human lncRNAs. A few of these novel lncRNAs were expressed in the rNETs, pNETs or siNETs. Further, computationally predicted regulatory mechanisms for a set of novel lncRNAs revealed direct correlation with cancer hallmark genes and molecular mechanisms of carcinogenesis and progression, which help understand the role of lncRNAs in neuroendocrine tumors. Our analyses provide the base for further experimental validations of these interactions. Data reported here open up the field of lncRNA-based regulation in GI-NETs.

## Abbreviations

DEMis: Differentially expressed miRNAs
GI-NETs: Gastrointestinal neuroendocrine tumors
GO: Gene ontology
lincRNA: long intergenic noncoding RNA
lncRNAs: Long non-coding RNAs
NET: Neuroendocrine tumors
PCGs: Protein-coding genes
PCTs: Protein-coding transcripts
pNETs: Pancreatic neuroendocrine tumors
rNETS: Rectal neuroendocrine tumors
siNETs: Small interstine neuroendocrine tumors

## Declarations

### Funding

The research work presented in this manuscript did not receive any specific grant from public, commercial, or not-for-profit funding agencies.

### Ethical approval and consent to participate

Not applicable

### Competing interests

The authors declare that they do not have competing interest.

## Authors contribution

Mahesh Kumar Padwal: Conceptualization; Data curation; Formal analysis; Investigation; Methodology; Software; Visualization; Roles/Writing - original draft.

Bhakti Basu: Conceptualization; Project administration; Validation; Resources; Supervision; Roles/Writing - review & editing.

## Acknowledgements

Not applicable.

## Availability of the data

This study has utilized the data from the public repository. The accession number and details of data are appropriately cited. The data generated from the current study are available in supplementary files with the manuscript submission.

## Code Avaibility

Pipeline and Code used for the analysis can be make avavilble to reviewers’s for evalution purpose on requests.

## Consent to participate

All authors have their consent to participate.

## Consent for publication

All authors have their consent to publish this article.

## Legends to Supplementary Figures

**Supplementary Figure S1:**
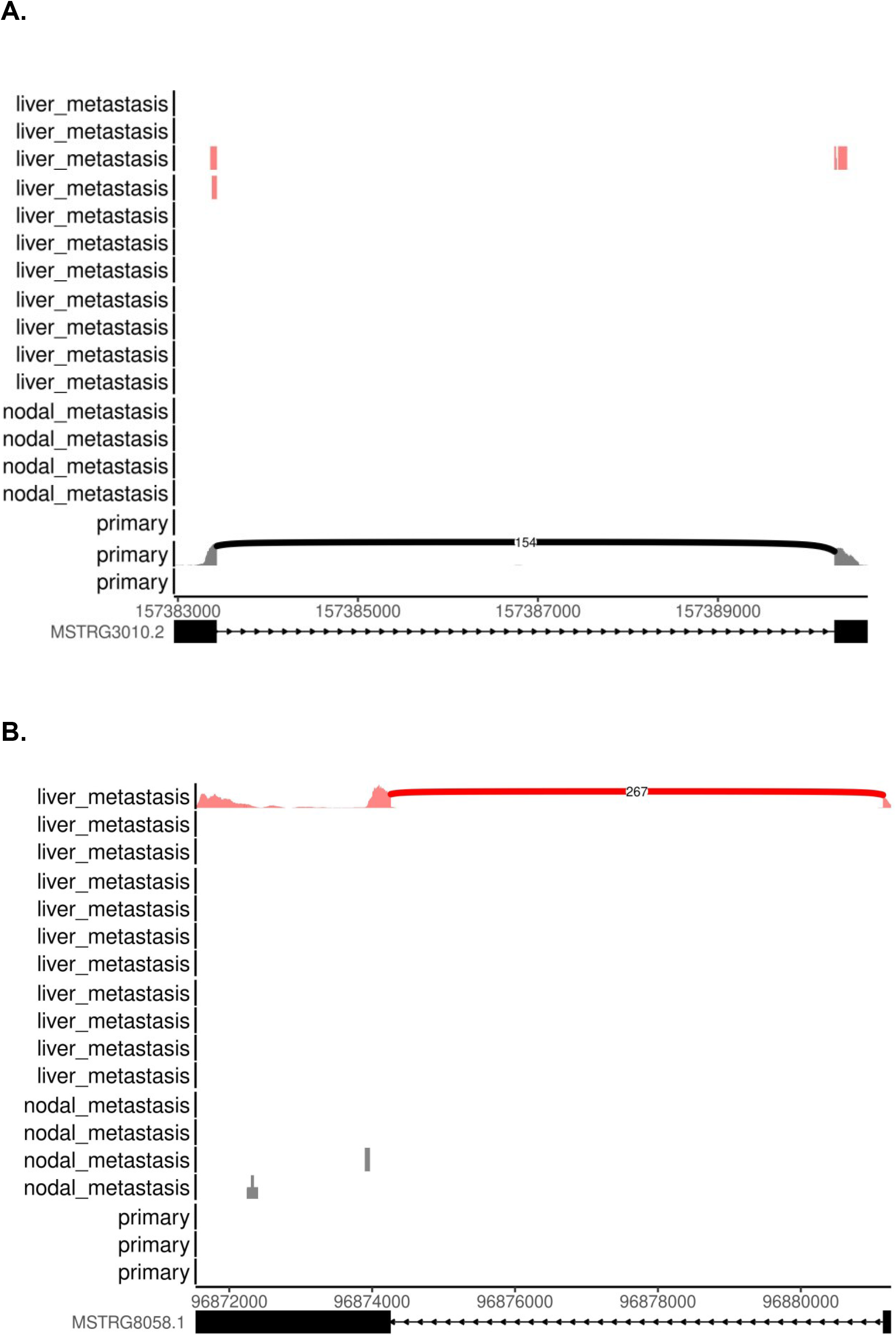
Sashimi plot for the lncRNA MSTRG3010.2 (A) and MSTRG8058.1 (B) are shown. Numbers on the x-axis indicate the coordinates of the transcript on chromosome and numbers along the transcripts indicate exon coverage.

**Supplementary Figure S2:**
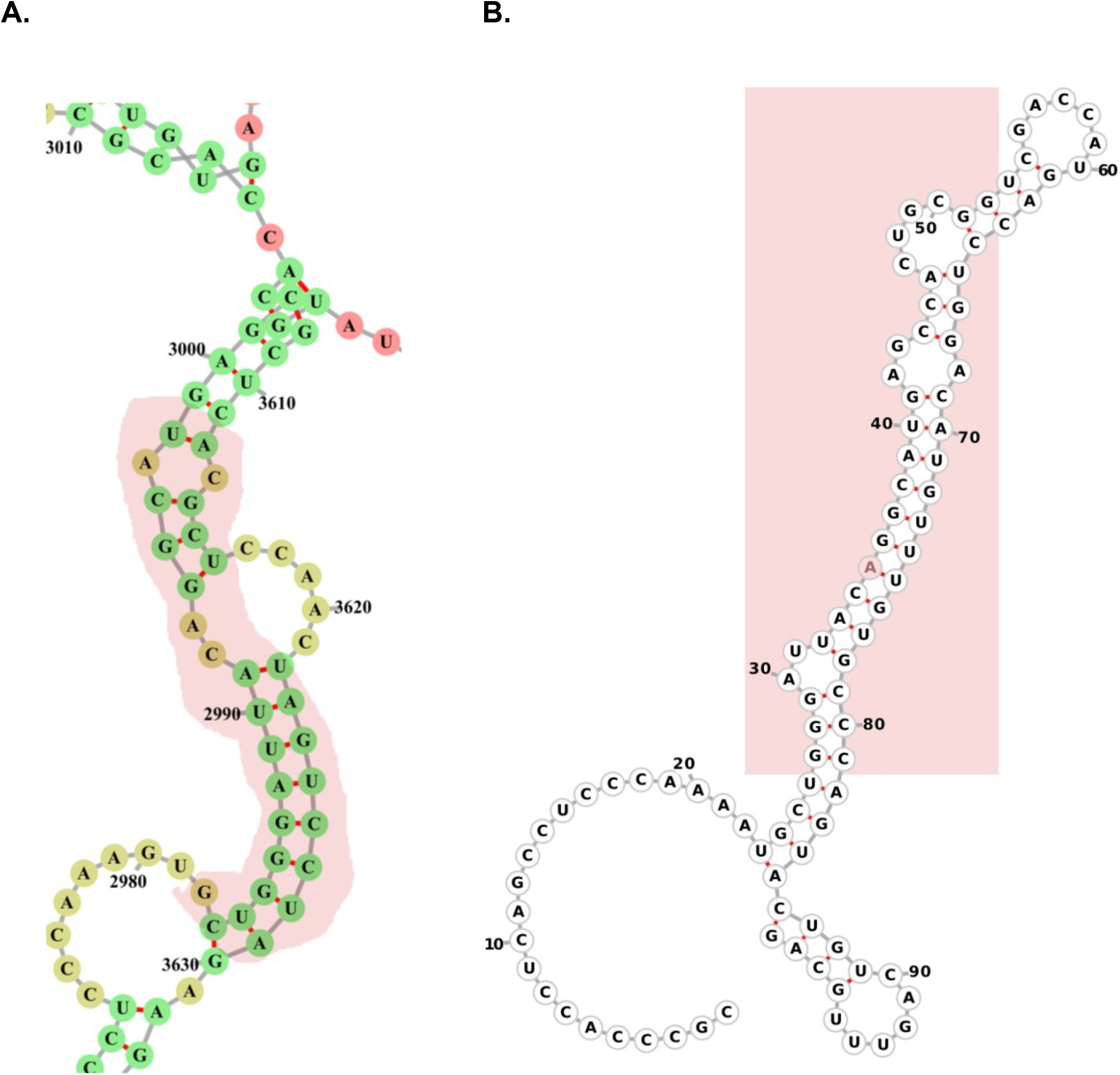
Hairpin stem-loop structure of the lncRNA MSTRG2354.2 (A) and precursors miRNA hsa-miR-619 (B) containing the mature miRNA sequence. The mature miRNA sequence is highlighted in the pink color.

**Supplementary Figure S3:**
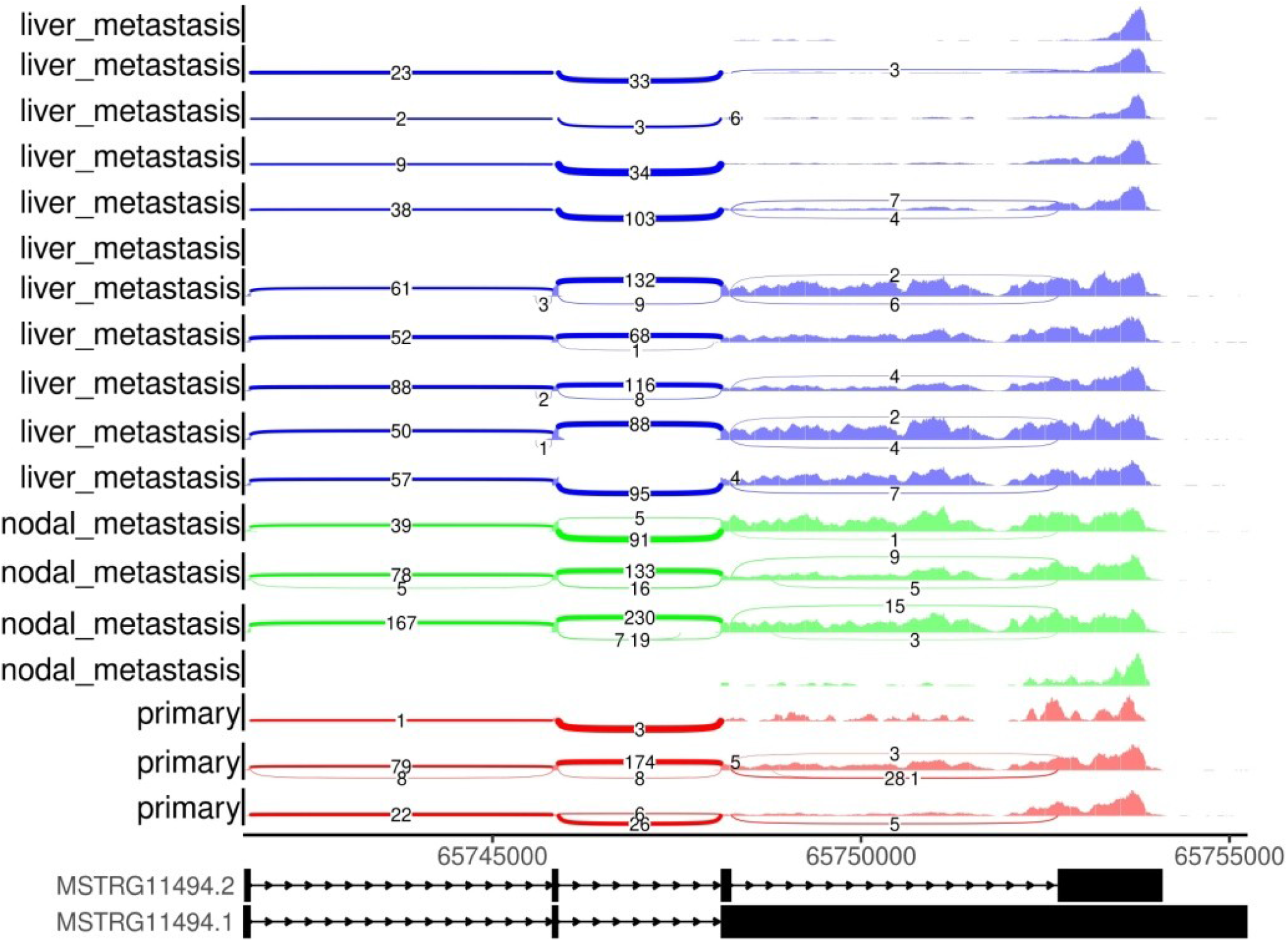
Sashimi plot for the lncRNA MSTRG11494.1. Numbers on the x-axis indicate the coordinates of the transcript on chromosome and numbers along the transcripts indicate exon coverage.

## Legends to Tables

**Table 1:** Read statistics of the downloaded SRA Libraries.

**Table 2:** mechRNA predicted regulatory mechanisms for important lncRNAs for cancer hallmarks genes.

## Legends to Supplementary Tables

**Table S1:** Number of the *de novo* assembled transcripts in each RNA-SEQ library.

**Table S2:** Chromosomal distribution of the identified novel lncRNAs.

**Table S3:** Phastcons conservation score for the identified novel lncRNA transcripts.

**Table S4:** Highly expressed novel lncRNAs in rNETs.

**Table S5:** TPM counts for the novel lncRNAs in normal rectal mucosa samples.

**Table S6:** TPM counts for the novel lncRNAs in pNETs.

**Table S7:** TPM counts for the novel lncRNAs in siNETs.

**Table S8:** Predicted probable location of the novel lncRNAs in one of the four compartments using LocateR.

**Table S9:** List of the hsa-miRNAs targeting novel lncRNAs.

**Table S10:** List of the lncRNAs having multiple binding sites (> 2) for miRNA.

**Table S11:** List of the lncRNAs targeted by DEMis reported to be important in differentiating between normal rectal mucosa and rectal carcinoma.

**Table S12:** List of PCGs predicted to be cis-regulated by novel lncRNAs.

**Table S13:** List of PCGs located within 100 kb region of the lncRNAs.

**Table S14:** List of PCGs located within 10 kb region of the lncRNA.

**Table S15:** List of the bi-directional lncRNAs located within 1 kb region from PCGs.

**Table S16:** Gene ontology terms significantly associated with *cis*-PCGs.

**Table S17:** Significantly enriched Reactome pathways associated with *cis*-PCGs.

**Table S18:** List of PCGs predicted to be trans-regulated by novel lncRNAs and their correlation coefficient.

**Table S19:** Significantly enriched Reactome pathways associated with *trans*-PCGs.

**Table S20:** Gene ontology terms significantly associated with *trans*-PCGs.

**Table S21:** List of the PCGs associated with novel lncRNA MSTRG16489.

**Table S22:** List of the RBPs associated with the novel lncRNAs.

**Table S23:** mechRNA mediated predicted mechanisms for important DEMs and lncRNAs.

## Legends to Additional Files

**Additional File 1:** Parameters used for aligning reads to the genome using STAR aligner.

**Additional File 2**: A sequences of the novel identified lncRNAs in fasta format.

**Additional File 3**: GTF file format for the novel identified lncRNAs

## Notes

### Competing Interest Statement

The authors have declared no competing interest.

## References

Ahmad A, Lin H, Shatabda S (2020) Locate-R: Subcellular localization of long non-coding RNAs using nucleotide compositions. Genomics 112:2583–2589. DOI 10.1016/j.ygeno.2020.02.011

Alexander RP, Fang G, Rozowsky J, Snyder M, Gerstein MB (2010) Annotating non-coding regions of the genome. Nat Rev Genet 11:559–571. DOI 10.1038/nrg2814

Alvarez MJ, Subramaniam PS, Tang LH, Grunn A, Aburi M, Rieckhof G, Komissarova EV, Hagan EA, Bodei L, Clemons PA, Dela Cruz FS, Dhall D, Diolaiti D, Fraker DA, Ghavami A, Kaemmerer D, Karan C, Kidd M, Kim KM, Kim HC, Kunju LP, Langel U, Li Z, Lee J, Li H, LiVolsi V, Pfragner R, Rainey AR, Realubit RB, Remotti H, Regberg J, Roses R, Rustgi A, Sepulveda AR, Serra S, Shi C, Yuan X, Barberis M, Bergamaschi R, Chinnaiyan AM, Detre T, Ezzat S, Frilling A, Hommann M, Jaeger D, Kim MK, Knudsen BS, Kung AL, Leahy E, Metz DC, Milsom JW, Park YS, Reidy-Lagunes D, Schreiber S, Washington K, Wiedenmann B, Modlin I, Califano A (2018) A precision oncology approach to the pharmacological targeting of mechanistic dependencies in neuroendocrine tumors. Nat Genet 50:979–989. DOI 10.1038/s41588-018-0138-4

Amodio N, Raimondi L, Juli G, Stamato MA, Caracciolo D, Tagliaferri P, Tassone P (2018) MALAT1: a druggable long non-coding RNA for targeted anti-cancer approaches. J Hematol Oncol 11:63. DOI 10.1186/s13045-018-0606-4

Bergenfelz C, Gaber A, Allaoui R, Mehmeti M, Jirstrom K, Leanderson T, Leandersson K (2015) S100A9 expressed in ER(-)PgR(-) breast cancers induces inflammatory cytokines and is associated with an impaired overall survival. Br J Cancer 113:1234–1243. DOI 10.1038/bjc.2015.346

Bolger AM, Lohse M, Usadel B (2014) Trimmomatic: a flexible trimmer for Illumina sequence data. Bioinformatics 30:2114–2120. DOI 10.1093/bioinformatics/btu170

Briata P, Gherzi R (2020) Long Non-Coding RNA-Ribonucleoprotein Networks in the Post-Transcriptional Control of Gene Expression. Non-coding RNA 6. DOI 10.3390/ncrna6030040

Cao H, Wahlestedt C, Kapranov P (2018) Strategies to Annotate and Characterize Long Noncoding RNAs: Advantages and Pitfalls. Trends Genet 34:704–721. DOI 10.1016/j.tig.2018.06.002

Chen J, Ju HL, Yuan XY, Wang TJ, Lai BQ (2016) SOX4 is a potential prognostic factor in human cancers: a systematic review and meta-analysis. Clinical & translational oncology : official publication of the Federation of Spanish Oncology Societies and of the National Cancer Institute of Mexico 18:65–72. DOI 10.1007/s12094-015-1337-4

Chiu HS, Somvanshi S, Patel E, Chen TW, Singh VP, Zorman B, Patil SL, Pan Y, Chatterjee SS, Cancer Genome Atlas Research N, Sood AK, Gunaratne PH, Sumazin P (2018) Pan-Cancer Analysis of lncRNA Regulation Supports Their Targeting of Cancer Genes in Each Tumor Context. Cell Rep 23:297–312 e212. DOI 10.1016/j.celrep.2018.03.064

Cong Z, Diao Y, Xu Y, Li X, Jiang Z, Shao C, Ji S, Shen Y, De W, Qiang Y (2019) Long non-coding RNA linc00665 promotes lung adenocarcinoma progression and functions as ceRNA to regulate AKR1B10-ERK signaling by sponging miR-98. Cell Death Dis 10:84. DOI 10.1038/s41419-019-1361-3

Djebali S, Davis CA, Merkel A, Dobin A, Lassmann T, Mortazavi A, Tanzer A, Lagarde J, Lin W, Schlesinger F, Xue C, Marinov GK, Khatun J, Williams BA, Zaleski C, Rozowsky J, Roder M, Kokocinski F, Abdelhamid RF, Alioto T, Antoshechkin I, Baer MT, Bar NS, Batut P, Bell K, Bell I, Chakrabortty S, Chen X, Chrast J, Curado J, Derrien T, Drenkow J, Dumais E, Dumais J, Duttagupta R, Falconnet E, Fastuca M, Fejes-Toth K, Ferreira P, Foissac S, Fullwood MJ, Gao H, Gonzalez D, Gordon A, Gunawardena H, Howald C, Jha S, Johnson R, Kapranov P, King B, Kingswood C, Luo OJ, Park E, Persaud K, Preall JB, Ribeca P, Risk B, Robyr D, Sammeth M, Schaffer L, See LH, Shahab A, Skancke J, Suzuki AM, Takahashi H, Tilgner H, Trout D, Walters N, Wang H, Wrobel J, Yu Y, Ruan X, Hayashizaki Y, Harrow J, Gerstein M, Hubbard T, Reymond A, Antonarakis SE, Hannon G, Giddings MC, Ruan Y, Wold B, Carninci P, Guigo R, Gingeras TR (2012) Landscape of transcription in human cells. Nature 489:101–108. DOI 10.1038/nature11233

Dobin A, Davis CA, Schlesinger F, Drenkow J, Zaleski C, Jha S, Batut P, Chaisson M, Gingeras TR (2013) STAR: ultrafast universal RNA-seq aligner. Bioinformatics 29:15–21. DOI 10.1093/bioinformatics/bts635

Dozmorov MG, Giles CB, Koelsch KA, Wren JD (2013) Systematic classification of non-coding RNAs by epigenomic similarity. BMC Bioinformatics 14 Suppl 14:S2. DOI 10.1186/1471-2105-14-S14-S2

Eddy SR (2011) Accelerated Profile HMM Searches. PLoS Comput Biol 7:e1002195. DOI 10.1371/journal.pcbi.1002195

Fang J, Xiao L, Zhang Q, Peng Y, Wang Z, Liu Y (2020) Junction plakoglobin, a potential prognostic marker of oral squamous cell carcinoma, promotes proliferation, migration and invasion. Journal of oral pathology & medicine : official publication of the International Association of Oral Pathologists and the American Academy of Oral Pathology 49:30–38. DOI 10.1111/jop.12952

Fatima R, Akhade VS, Pal D, Rao SM (2015) Long noncoding RNAs in development and cancer: potential biomarkers and therapeutic targets. Mol Cell Ther 3:5. DOI 10.1186/s40591-015-0042-6

Garrido-Martin D, Palumbo E, Guigo R, Breschi A (2018) ggsashimi: Sashimi plot revised for browser- and annotation-independent splicing visualization. PLoS Comput Biol 14:e1006360. DOI 10.1371/journal.pcbi.1006360

Gawronski AR, Uhl M, Zhang Y, Lin YY, Niknafs YS, Ramnarine VR, Malik R, Feng F, Chinnaiyan AM, Collins CC, Sahinalp SC, Backofen R (2018) MechRNA: prediction of lncRNA mechanisms from RNA-RNA and RNA-protein interactions. Bioinformatics 34:3101–3110. DOI 10.1093/bioinformatics/bty208

Grebhardt S, Muller-Decker K, Bestvater F, Hershfinkel M, Mayer D (2014) Impact of S100A8/A9 expression on prostate cancer progression in vitro and in vivo. J Cell Physiol 229:661–671. DOI 10.1002/jcp.24489

Gyvyte U, Kupcinskas J, Juzenas S, Inciuraite R, Poskiene L, Salteniene V, Link A, Fassan M, Franke A, Kupcinskas L, Skieceviciene J (2018) Identification of long intergenic non-coding RNAs (lincRNAs) deregulated in gastrointestinal stromal tumors (GISTs). PLoS One 13:e0209342. DOI 10.1371/journal.pone.0209342

Haase D, Meister M, Muley T, Hess J, Teurich S, Schnabel P, Hartenstein B, Angel P (2007) FRMD3, a novel putative tumour suppressor in NSCLC. Oncogene 26:4464–4468. DOI 10.1038/sj.onc.1210225

Han J, Meng J, Chen S, Wang X, Yin S, Zhang Q, Liu H, Qin R, Li Z, Zhong W, Zhang C, Zhang H, Tang Y, Lin T, Gao W, Zhang X, Yang L, Liu Y, Zhou HG, Sun T, Yang C (2019) YY1 Complex Promotes Quaking Expression via Super-Enhancer Binding during EMT of Hepatocellular Carcinoma. Cancer research 79:1451–1464. DOI 10.1158/0008-5472.CAN-18-2238

Han Li C, Chen Y (2015) Small and Long Non-Coding RNAs: Novel Targets in Perspective Cancer Therapy. Curr Genomics 16:319–326. DOI 10.2174/1389202916666150707155851

Han X, Ji Y, Zhao J, Xu X, Lou W (2013) Expression of PTEN and mTOR in pancreatic neuroendocrine tumors. Tumour Biol 34:2871–2879. DOI 10.1007/s13277-013-0849-1

Hesson LB, Ng B, Zarzour P, Srivastava S, Kwok CT, Packham D, Nunez AC, Beck D, Ryan R, Dower A, Ford CE, Pimanda JE, Sloane MA, Hawkins NJ, Bourke MJ, Wong JW, Ward RL (2016) Integrated Genetic, Epigenetic, and Transcriptional Profiling Identifies Molecular Pathways in the Development of Laterally Spreading Tumors. Mol Cancer Res 14:1217–1228. DOI 10.1158/1541-7786.MCR-16-0175

Higurashi M, Iketani M, Takei K, Yamashita N, Aoki R, Kawahara N, Goshima Y (2012) Localized role of CRMP1 and CRMP2 in neurite outgrowth and growth cone steering. Developmental neurobiology 72:1528–1540. DOI 10.1002/dneu.22017

Huang HY, Lin YC, Li J, Huang KY, Shrestha S, Hong HC, Tang Y, Chen YG, Jin CN, Yu Y, Xu JT, Li YM, Cai XX, Zhou ZY, Chen XH, Pei YY, Hu L, Su JJ, Cui SD, Wang F, Xie YY, Ding SY, Luo MF, Chou CH, Chang NW, Chen KW, Cheng YH, Wan XH, Hsu WL, Lee TY, Wei FX, Huang HD (2020) miRTarBase 2020: updates to the experimentally validated microRNA-target interaction database. Nucleic Acids Res 48:D148–D154. DOI 10.1093/nar/gkz896

Jeong H, Kim J, Lee Y, Seo JH, Hong SR, Kim A (2014) Neuregulin-1 induces cancer stem cell characteristics in breast cancer cell lines. Oncology reports 32:1218–1224. DOI 10.3892/or.2014.3330

John B, Enright AJ, Aravin A, Tuschl T, Sander C, Marks DS (2004) Human MicroRNA targets. PLoS Biol 2:e363. DOI 10.1371/journal.pbio.0020363

Juang YL, Jeng YM, Chen CL, Lien HC (2016) PRRX2 as a novel TGF-beta-induced factor enhances invasion and migration in mammary epithelial cell and correlates with poor prognosis in breast cancer. Molecular carcinogenesis 55:2247–2259. DOI 10.1002/mc.22465

Kaji H, Canaff L, Lebrun JJ, Goltzman D, Hendy GN (2001) Inactivation of menin, a Smad3-interacting protein, blocks transforming growth factor type beta signaling. Proc Natl Acad Sci U S A 98:3837–3842. DOI 10.1073/pnas.061358098

Kornienko AE, Dotter CP, Guenzl PM, Gisslinger H, Gisslinger B, Cleary C, Kralovics R, Pauler FM, Barlow DP (2016) Long non-coding RNAs display higher natural expression variation than protein-coding genes in healthy humans. Genome Biol 17:14. DOI 10.1186/s13059-016-0873-8

Lee JH, Hur W, Hong SW, Kim JH, Kim SM, Lee EB, Yoon SK (2017) ELK3 promotes the migration and invasion of liver cancer stem cells by targeting HIF-1alpha. Oncology reports 37:813–822. DOI 10.3892/or.2016.5293

Li A, Zhang J, Zhou Z (2014) PLEK: a tool for predicting long non-coding RNAs and messenger RNAs based on an improved k-mer scheme. BMC Bioinformatics 15:311. DOI 10.1186/1471-2105-15-311

Li SC, Essaghir A, Martijn C, Lloyd RV, Demoulin JB, Oberg K, Giandomenico V (2013) Global microRNA profiling of well-differentiated small intestinal neuroendocrine tumors. Mod Pathol 26:685–696. DOI 10.1038/modpathol.2012.216

Man D, Wu J, Shen Z, Zhu X (2018) Prognosis of patients with neuroendocrine tumor: a SEER database analysis. Cancer Manag Res 10:5629–5638. DOI 10.2147/CMAR.S174907

Marusiak AA, Prelowska MK, Mehlich D, Lazniewski M, Kaminska K, Gorczynski A, Korwat A, Sokolowska O, Kedzierska H, Golab J, Biernat W, Plewczynski D, Brognard J, Nowis D (2019) Upregulation of MLK4 promotes migratory and invasive potential of breast cancer cells. Oncogene 38:2860–2875. DOI 10.1038/s41388-018-0618-0

Nagy A, Munkacsy G, Gyorffy B (2021) Pancancer survival analysis of cancer hallmark genes. Scientific reports 11:6047. DOI 10.1038/s41598-021-84787-5

Neph S, Kuehn MS, Reynolds AP, Haugen E, Thurman RE, Johnson AK, Rynes E, Maurano MT, Vierstra J, Thomas S, Sandstrom R, Humbert R, Stamatoyannopoulos JA (2012) BEDOPS: high-performance genomic feature operations. Bioinformatics 28:1919–1920. DOI 10.1093/bioinformatics/bts277

Ohno S (1972) So much “junk” DNA in our genome. Brookhaven Symp Biol 23:366–370

Papageorgis P, Cheng K, Ozturk S, Gong Y, Lambert AW, Abdolmaleky HM, Zhou JR, Thiagalingam S (2011) Smad4 inactivation promotes malignancy and drug resistance of colon cancer. Cancer research 71:998–1008. DOI 10.1158/0008-5472.CAN-09-3269

Pellatt DF, Stevens JR, Wolff RK, Mullany LE, Herrick JS, Samowitz W, Slattery ML (2016) Expression Profiles of miRNA Subsets Distinguish Human Colorectal Carcinoma and Normal Colonic Mucosa. Clin Transl Gastroenterol 7:e152. DOI 10.1038/ctg.2016.11

Pertea G, Pertea M (2020) GFF Utilities: GffRead and GffCompare. F1000Res 9:304. DOI 10.12688/f1000research.23297.1

Pertea M (2012) The human transcriptome: an unfinished story. Genes (Basel) 3:344–360. DOI 10.3390/genes3030344

Pertea M, Pertea GM, Antonescu CM, Chang TC, Mendell JT, Salzberg SL (2015) StringTie enables improved reconstruction of a transcriptome from RNA-seq reads. Nat Biotechnol 33:290–295. DOI 10.1038/nbt.3122

Prensner JR, Iyer MK, Balbin OA, Dhanasekaran SM, Cao Q, Brenner JC, Laxman B, Asangani IA, Grasso CS, Kominsky HD, Cao X, Jing X, Wang X, Siddiqui J, Wei JT, Robinson D, Iyer HK, Palanisamy N, Maher CA, Chinnaiyan AM (2011) Transcriptome sequencing across a prostate cancer cohort identifies PCAT-1, an unannotated lincRNA implicated in disease progression. Nat Biotechnol 29:742–749. DOI 10.1038/nbt.1914

Rindi G, Wiedenmann B (2011) Neuroendocrine neoplasms of the gut and pancreas: new insights. Nat Rev Endocrinol 8:54–64. DOI 10.1038/nrendo.2011.120

Rodrigues A, Castro-Pocas F, Pedroto I (2015) Neuroendocrine Rectal Tumors: Main Features and Management. GE Port J Gastroenterol 22:213–220. DOI 10.1016/j.jpge.2015.04.008

Romano G, Veneziano D, Acunzo M, Croce CM (2017) Small non-coding RNA and cancer. Carcinogenesis 38:485–491. DOI 10.1093/carcin/bgx026

Sanchez Calle A, Kawamura Y, Yamamoto Y, Takeshita F, Ochiya T (2018) Emerging roles of long non-coding RNA in cancer. Cancer Sci 109:2093–2100. DOI 10.1111/cas.13642

Sato F, Bhawal UK, Yoshimura T, Muragaki Y (2016) DEC1 and DEC2 Crosstalk between Circadian Rhythm and Tumor Progression. J Cancer 7:153–159. DOI 10.7150/jca.13748

Srikrishna G (2012) S100A8 and S100A9: new insights into their roles in malignancy. J Innate Immun 4:31–40. DOI 10.1159/000330095

Tan F, Thiele CJ, Li Z (2014) Collapsin response mediator proteins: Potential diagnostic and prognostic biomarkers in cancers (Review). Oncology letters 7:1333–1340. DOI 10.3892/ol.2014.1909

Volders PJ, Anckaert J, Verheggen K, Nuytens J, Martens L, Mestdagh P, Vandesompele J (2019) LNCipedia 5: towards a reference set of human long non-coding RNAs. Nucleic Acids Res 47:D135–D139. DOI 10.1093/nar/gky1031

Wang L, Cho KB, Li Y, Tao G, Xie Z, Guo B (2019) Long Noncoding RNA (lncRNA)-Mediated Competing Endogenous RNA Networks Provide Novel Potential Biomarkers and Therapeutic Targets for Colorectal Cancer. Int J Mol Sci 20. DOI 10.3390/ijms20225758

Wang L, Park HJ, Dasari S, Wang S, Kocher JP, Li W (2013) CPAT: Coding-Potential Assessment Tool using an alignment-free logistic regression model. Nucleic Acids Res 41:e74. DOI 10.1093/nar/gkt006

Watson CN, Belli A, Di Pietro V (2019) Small Non-coding RNAs: New Class of Biomarkers and Potential Therapeutic Targets in Neurodegenerative Disease. Front Genet 10:364. DOI 10.3389/fgene.2019.00364

Williams CB, Phelps-Polirer K, Dingle IP, Williams CJ, Rhett MJ, Eblen ST, Armeson K, Hill EG, Yeh ES (2020) HUNK phosphorylates EGFR to regulate breast cancer metastasis. Oncogene 39:1112–1124. DOI 10.1038/s41388-019-1046-5

Xu K, Qiu C, Pei H, Mehmood MA, Wang H, Li L, Xia Q (2018) Homeobox B3 promotes tumor cell proliferation and invasion in glioblastoma. Oncology letters 15:3712–3718. DOI 10.3892/ol.2018.7750

Yan W, Wu K, Herman JG, Xu X, Yang Y, Dai G, Guo M (2018) Retinoic acid-induced 2 (RAI2) is a novel tumor suppressor, and promoter region methylation of RAI2 is a poor prognostic marker in colorectal cancer. Clin Epigenetics 10:69. DOI 10.1186/s13148-018-0501-4

Yu G, Wang LG, Han Y, He QY (2012) clusterProfiler: an R package for comparing biological themes among gene clusters. OMICS 16:284–287. DOI 10.1089/omi.2011.0118

Zhang Y, Pitchiaya S, Cieslik M, Niknafs YS, Tien JC, Hosono Y, Iyer MK, Yazdani S, Subramaniam S, Shukla SK, Jiang X, Wang L, Liu TY, Uhl M, Gawronski AR, Qiao Y, Xiao L, Dhanasekaran SM, Juckette KM, Kunju LP, Cao X, Patel U, Batish M, Shukla GC, Paulsen MT, Ljungman M, Jiang H, Mehra R, Backofen R, Sahinalp CS, Freier SM, Watt AT, Guo S, Wei JT, Feng FY, Malik R, Chinnaiyan AM (2018) Analysis of the androgen receptor-regulated lncRNA landscape identifies a role for ARLNC1 in prostate cancer progression. Nat Genet 50:814–824. DOI 10.1038/s41588-018-0120-1

Zhang Y, Xu L, Li A, Han X (2019) The roles of ZEB1 in tumorigenic progression and epigenetic modifications. Biomed Pharmacother 110:400–408. DOI 10.1016/j.biopha.2018.11.112

Zhou RS, Zhang EX, Sun QF, Ye ZJ, Liu JW, Zhou DH, Tang Y (2019) Integrated analysis of lncRNA-miRNA-mRNA ceRNA network in squamous cell carcinoma of tongue. BMC Cancer 19:779. DOI 10.1186/s12885-019-5983-8

